# Co-translational determination of quaternary structures in chaperone factories

**DOI:** 10.1101/2025.02.12.637890

**Authors:** Manon Philippe, Soha Salloum, Floric Slimani, Héloïse Chassé, Marie-Cécile Robert, Serge Urbach, Jacques Imbert, Martial Séveno, Simon Georges, Séverine Boulon, Céline Verheggen, Edouard Bertrand

**Affiliations:** IGH, Montpellier University and CNRS, Montpellier, France; équipe labellisée Ligue Nationale Contre le Cancer; IGF, Univ. Montpellier, CNRS, INSERM, Montpellier, France; BCM, Univ. Montpellier, CNRS, INSERM, Montpellier, France

**Keywords:** co-translational assembly, chaperone, translation factories

## Abstract

The HSP90/R2TP quaternary chaperone assembles key cellular machines, including the three nuclear RNA polymerases and many non-coding RNPs. Here, we characterized the RNA associated to R2TP and found that it binds many partners co-translationally. Its co-translational interactome further reveals many novel potential clients and identifies clients bound only co-translationally, only post-translationally, or both. For pairs of subunits assembling together and bound co-translationally by R2TP, only a marginal proportion of their mRNAs is co-localized and co-translated. Instead, the HSP90 and R2TP chaperones induce the formation of condensates accumulating client mRNAs and thus favoring co-translational interactions between chaperones and clients. The R2TP then cycles between co- and post-translational steps and this is regulated by ATP: it binds co-translationally in absence of ATP and becomes released from post-translational assembly intermediates by ATP hydrolysis. Assembly of protein complexes is thus initiated early by chaperones and this mechanism, dubbed co-translational chaperone channeling (cha-cha), substitutes for the rarity of co-localized/co-translated mRNAs.

## Introduction

Non-coding RNPs are complex macromolecular assemblies and studies performed on many systems have established that they require specific factors for their assembly. For instance, a large number of comprehensive studies performed in the last 20 years have revealed that ∼200 assembly factors are required to produce functional ribosomes ^1^. Likewise, despite their apparent simplicity, small RNPs like the spliceosomal snRNPs or the nucleolar snoRNPs also require a surprisingly large number of specialized assembly factors ^2^. These assembly factors are evolutionary conserved and their number frequently exceeds the set of components of the mature RNP particle, highlighting the generality and importance of this process ^2,3^. Assembly machineries also occur for stable protein complexes devoid of RNAs. For instance, the formation of RNA polymerases requires an assembly machinery that includes more than 14 factors and whose mechanism of action remains poorly understood ^4^. Interestingly, some of these assembly factors, like the HSP90/R2TP chaperone, are linked to the protein folding machinery. Indeed, quaternary protein folding can be seen as an extension of tertiary folding, with the main difference being that it occurs in *trans* and not in *cis*. Assembly factors have been shown to chaperone unassembled subunits, improve efficiency and specificity of the assembly process, transport subunits across the cell and perform quality controls ^1,2,3^.

The R2TP is a conserved HSP90 co-chaperone that has a unique and essential role in the cell as a generalist assembly chaperone that helps build multi-subunit complexes ^5,6,7,8^. Specifically, the R2TP promotes the biogenesis of: (i) small non-coding RNPs such as the box C/D and H/ACA^2,9,10^ the telomerase RNP ^11,12^; U4 and U5 spliceosomal snRNPs ^13,14^; and miRNPs ^15^; (ii) the three nuclear RNA polymerases ^4,16^; (iii) ciliary dyneins ^17^; (iv) complexes containing PIKKs (mTOR, ATM/ATR, DNA-PK, TRRAP, SMG1) ^18,19^; and (v) the Tuberous Sclerosis Complex (TSC) that controls mTOR activity ^20^. This list of client complexes is likely incomplete, as large-scale interaction assays have revealed additional R2TP partners ^21,22,23,24^. In agreement with this diversity of roles, the R2TP subunit RPAP3 is essential in mice, and its targeted deletion in the adult intestine leads to rapid disruption of tissue homeostasis because of a lack of assembly of key complexes such as RNA polymerase II ^25^.

The R2TP is highly conserved across evolution and it can be tracked to the root of the eukaryotic lineage ^26^. It is composed of a heterodimer of RPAP3:PIH1D1 associated with the AAA+ ATPases RUVBL1 and RUVBL2 ^23,27^,. In humans, the R2TP further associates with a set of prefoldins and prefoldin-like subunits to form the PAQosome ^8^. RUVBL1/RUVBL2 form hetero-hexameric and dodecameric rings and are believed to carry a chaperone activity on their own ^28,29,30^. Compared to other AAA+ ATPases, they have an extra domain called DII, which is located outside the ring and whose conformational changes are believed to play important roles in the assembly function of R2TP ^7,28,31,32^. Indeed, biochemical and structural data have shown that RUVBL1/RUVBL2 make ATP-dependent contacts with clients and cofactors ^7,33^. Because of their hexameric nature, it is thought that this enables them to hold various subunits in place during the assembly process, with nucleotides acting as a switch for client binding or release (^34^ for a review). Human RPAP3 possesses two TPR domains that bind HSP70 and HSP90 and a C-terminal domain that associates with RUVBL1/RUVBL2 at the opposite side of DII ^26,31,35,36^. RPAP3 forms a heterodimer with PIH1D1, which is important for client recognition and the regulation of the ATPase activities of RUVBL1/RUVBL2 and HSP70/90 during their chaperone cycle ^35,37,38^. The recognition of R2TP clients can occur in several ways. Some clients are recruited with the help of cofactors, such as the TTT complex for PIKKs ^39,40,41^; ZNHIT2 and ECD for U5 snRNP ^7,13,14^; and NOPCHAP1, NUFIP1, ZNHIT3 and ZNHIT6 for C/D snoRNPs ^9,24,42,43^. Others are directly recruited by R2TP, and this can occur via the PIH domain of PIH1D1, which contains a basic pocket that binds phosphorylated acidic peptides phosphorylated of the DpSDD/E consensus motif ^41,44^, or via RPAP3, as in the case of the Dicer co-factor TRBP ^15^ or for clients containing Armadillo repeats ^23^.

Recent studies suggested that the assembly of multi-subunit complexes is intimately coupled with translation ^45,46,47^. For instance, biochemical studies in yeast and mammals have shown that subunits of the same complex can bind each other co-translationally, sometimes reciprocally and other times in a directional manner ^45^. In mammals, detailed studies of the general transcription factor TFIID and the co-activators SAGA and ATAC have revealed hierarchical co-translational assembly pathways that mix co- and post-translational steps ^48,49,50^. For TFIID in particular, some subunits join others during their synthesis to create sub-modules, which assemble post-translationally with the largest TAF1 subunit during its translation ^49^. Importantly, the co-translational joining of one subunit with another can occur either by associating and co-translating the two polysomes together (called the co-co pathway), or by the transport of one fully translated subunit to its partner polysome (the co-post pathway). Current evidence suggests that the co-co assembly pathway may be common ^51,52^ although single RNA imaging has so far only revealed only low frequencies of mRNA co-localization ^47,52^, leaving the exact usage of the co-co pathway an open question. In the case of the co-post pathway, how fully translated subunits are transported for assembly is not well understood. In the case of the SMN complex that assembles the Sm ring on snRNAs, the newly translated Sm proteins remain on the ribosome until they are picked up by the pICln assembly chaperone, which brings them to SMN ^53^. Despite the different mechanisms observed, many mechanisms converge to the ribosome, indicating that it may function as a hub driving assembly.

Here, we analyzed co-translational events for chaperone-assisted assembly of macromolecular complexes. We show that the HSP90/R2TP assembly chaperone binds co-translationally to many of its clients and that its co-translational interactome reveals many new partners. The co-co pathway appears to be marginal and instead, the chaperone induces the formation of dedicated translation factories that accumulate client mRNAs and thus favor chaperone-client co-translational interactions. The R2TP thus channels the formation of macromolecular complexes starting from subunit synthesis, likely compensating for the scarcity of subunit mRNA co-localization.

## Results

### RUVBL1 immuno-precipitates client mRNAs in a translation-dependent manner

The HSP90/R2TP chaperone (Figure 1A) is specialized in the assembly of macromolecular complexes and recent evidence has linked this process to translation (^47^ for a review). We thus tested whether the R2TP can bind its client proteins while they are being translated. We performed a RIP-Seq experiment using a heterozygous HCT116 cell line having GFP tag fused at the C-terminus of one allele of RUVBL1. This cell line was characterized by PCR genotyping, Western blot and immunofluorescence, confirming the proper insertion of the tag and the localization of the RUVBL1-GFP fusion in the cytoplasm and the nucleoplasm as expected (Figure S1A-B and Figure 1B, ^54^). We then immunoprecipitated (IP’ed) RUVBL1-GFP using GFP-Trap beads and sequenced the co-precipitated RNAs. Remarkably, we found 172 RNAs specifically enriched over the control IP (FDR<0.05), with enrichment values varying from 84 to 2 fold (Figure 1C and Table S1). Some mRNAs coded for known substrates of R2TP, such as the PRPF8 and EFTUD2 proteins of U5 snRNP, the large subunits of the three nuclear RNA polymerases (POLR1A, POLR2A, POLR2B, POLR3A) and the PIKK SMG1. Other well-known R2TP clients such as the core snoRNP proteins DKC1 and NOP58 did not pass the statistical threshold but were nevertheless among the top enriched mRNAs (ranked 194 and 330, respectively). RUVBL1/RUVBL2 are part of several chromatin complexes (TIP60, SRCAP, INO80), and this is believed to be independent of their participation into the R2TP chaperone. Remarkably, the mRNAs coding for key subunits of these complexes were also highly enriched in the IP (EP400, SRCAP, BRD8, INO80). Most interestingly, many mRNAs coding for proteins not previously known to interact with RUVBL1 were also found, and several formed clearly identifiable families of related proteins. This was the case for the AGO proteins (AGO1, AGO3 and AGO4, with also AGO2 in the top hits although below the statistical threshold), the CREB3 family (CREBRF, CREB3 and CREB3L2) and the SCAF family that share a similar RNA polymerase II CTD interacting domain (PCF11, SCAF4, SCAF8, CHERP). It was also noteworthy that RUVBL1-GFP associated with mRNAs coding for pairs of subunits that assemble together in mature complexes, such as POLR2A and POLR2B in RNA polymerase II, PRPF8 and EFTUD2 in U5 snRNP, and BRD8 and EP400 in the TIP60 chromatin remodeling complex. Other mRNAs coded for seemingly unrelated proteins, although many were long proteins, larger than 1000 amino acids. Finally, we also found an enrichment for several box C/D snoRNAs (SNORD3, SNORD13, SNORD46 and SNORD31), as expected from previous studies ^54,55^. In order to determine if the association of RUVBL1 with mRNAs was translation-dependent, we repeated the RIP-Seq experiment after a 30 min treatment with puromycin, which acts as a translation terminator and releases the nascent peptides from the ribosome. We found that most of the mRNAs previously enriched in the RUVBL1-GFP IP were lost or considerably less enriched (Figure 1D and S1C), indicating that RUVBL1-GFP binds to polysomes via the nascent proteins that are being translated.

**Figure 1.**
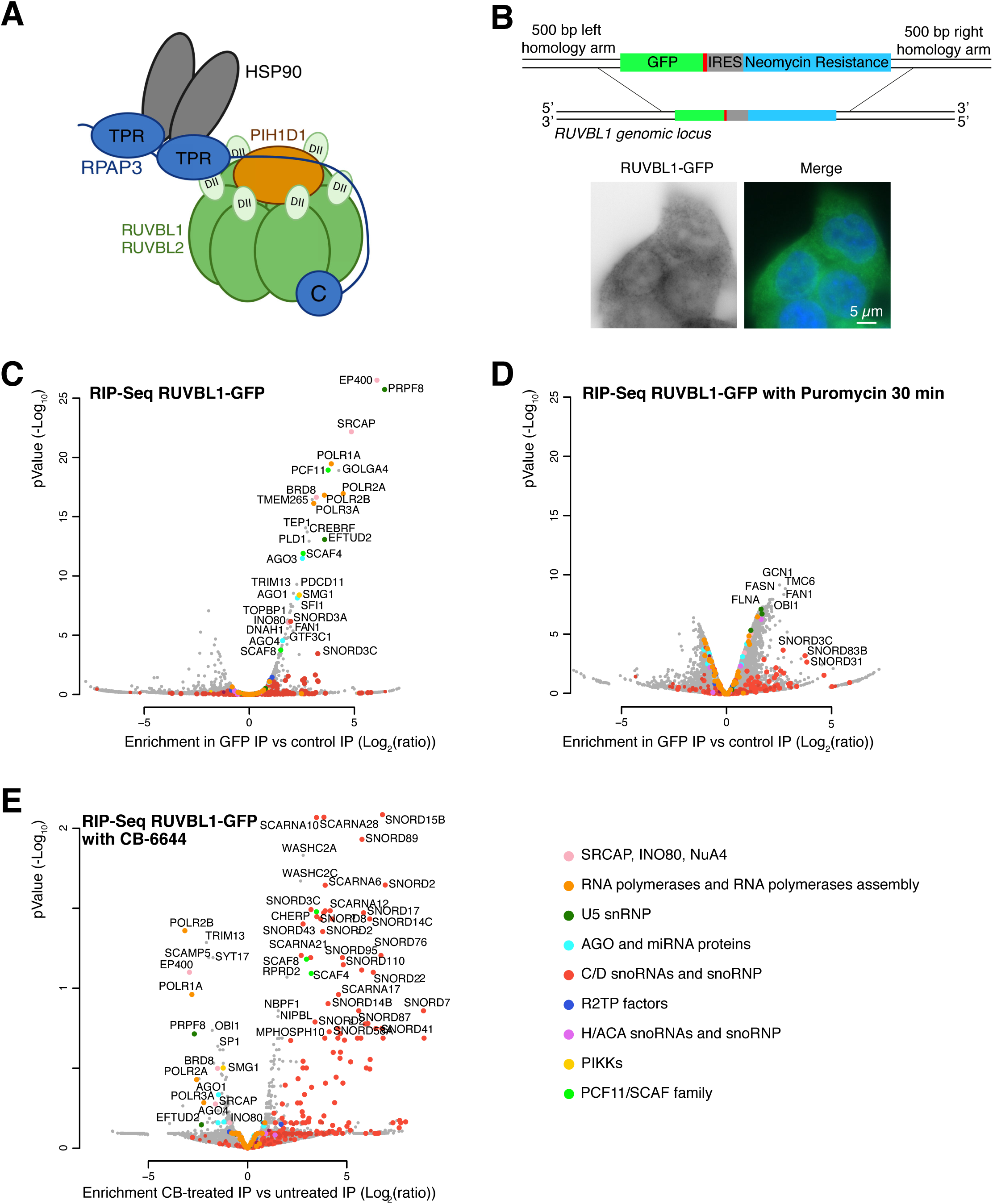
RUVBL1 associates co-translationaly with mRNAs. **A-** Schematic of the HSP90/R2TP chaperone. DII is the domain II of RUVBL1 and RUVBL2. **B-** Schematic of the cassette integrated at the RUVBL1 genomic locus and micrographs of HCT116 cells with a RUVBL1-GFP allele and imaged by fluorescent microscopy. Green: RUVBL1-GFP. Blue: DAPI staining. Scale bar: 5 µm. **C-** Volcano plot of the RUVBL1-GFP RIP-Seq experiment, showing the pValue (y axis; -Log10) as a function of the enrichment in the GFP IP over the control IP (x axis; Log2(ratio)). Each dot represents an mRNA that is colored according to the code on the right of panel E. **D-** Volcano plot of the RUVBL1-GFP RIP-Seq experiment in puromycin treated cells (Legend as in C). **E-** Vulcano plot comparing the RUVBL1-GFP RIP-seq in untreated versus CB-6644 treated condition. The graph shows the pValue (y axis; -Log10) as a function of the fold change in the GFP IPs of treated versus non-treated cells (x axis; Log2(ratio)).

### The ATPase activity of RUVBL1/RUVBL2 regulates their binding to polysomes

The AAA+ ATPase activity of RUVBL1/RUVBL2 is central to their chaperone activity and the nucleotide bound state (ATP, ADP or none) controls which partners they interact with ^28,31,33,55^. To test how ATP affects their interactions with RNAs, we performed a RIP-seq experiment in presence of CB-6644, a small molecule that inhibits their ATPase activity and blocks them in the ATP-bound state ^56,57^). Most of the mRNAs previously found in the RUVBL1-GFP IP were lost or less enriched after CB-6644 treatment, including the mRNAs coding for the U5 snRNP proteins and the subunits of the RNA polymerases and the chromatin remodeling complexes (Figure 1E and S1D). RUVBL1 may thus bind the corresponding nascent proteins in the ADP or empty state. In contrast, many box C/D snoRNAs were associated more strongly with RUVBL1-GFP upon CB-6644 treatment, suggesting that the drug prevents the release of RUVBL1/RUVBL2 from immature C/D snoRNPs. A few mRNAs also displayed stronger binding with CB-6644. This was the case for the PCF11/SCAF family (with RPRD2 as an additional member), and for WASHC2A and WAHSC2C mRNAs, which encode two almost identical proteins involved in actin dynamics. RUVBL1/RUVBL2 may thus bind more strongly to these nascent proteins when loaded with ATP. Altogether, this indicated that ATP regulates the co-translational binding of RUVBL1.

To gain more insight into this mechanism, we performed a quantitative label-free proteomic analysis of the RUVBL1-GFP IPs (AP-MS), with and without CB-6644 treatment. The proteomic analysis revealed the known partners of RUVBL1 as expected, including all subunits of the R2TP, the PAQosome and the INO80, SRCAP and TIP60 chromatin remodeling complexes, as well as many subunits of the three nuclear RNA polymerases, the U5 snRNP and the C/D and H/ACA snoRNPs (Figure 2A, Table S2). RUVBL1-GFP was also associated with known R2TP co-factors, such as ZNHIT2, ZNHIT3, NUFIP1, ZNHIT6, NOPCHAP1 and the TTT complex. In the presence of CB-6644, the subunits of the chromatin remodeling complexes became less enriched, while most of the other clients, such as the H/ACA core component DKC1 or the RNA polymerase II subunits, became more enriched (Figure 2B). As in the RIP-seq, binding to WAHSC2A increased dramatically after CB-6644 treatment. This was also the case for the C/D core component SNU13, which directly binds snoRNAs and likely reflects the increased binding of RUVBL1-GFP to C/D snoRNAs observed in the RIP-seq with CB-6644 (Figure 1E). Some R2TP cofactors showed increased binding, such as the TTT complex, while others dissociated (NOPCHAP1, DPCD, WDCP). This was consistent with the ATP-dependence observed *in vitro* for these proteins ^42,58^, suggesting that the differences observed *in vivo* reflect the genuine ATP-dependent affinity of RUVBL1/RUVBL2 for these partners.

**Figure 2.**
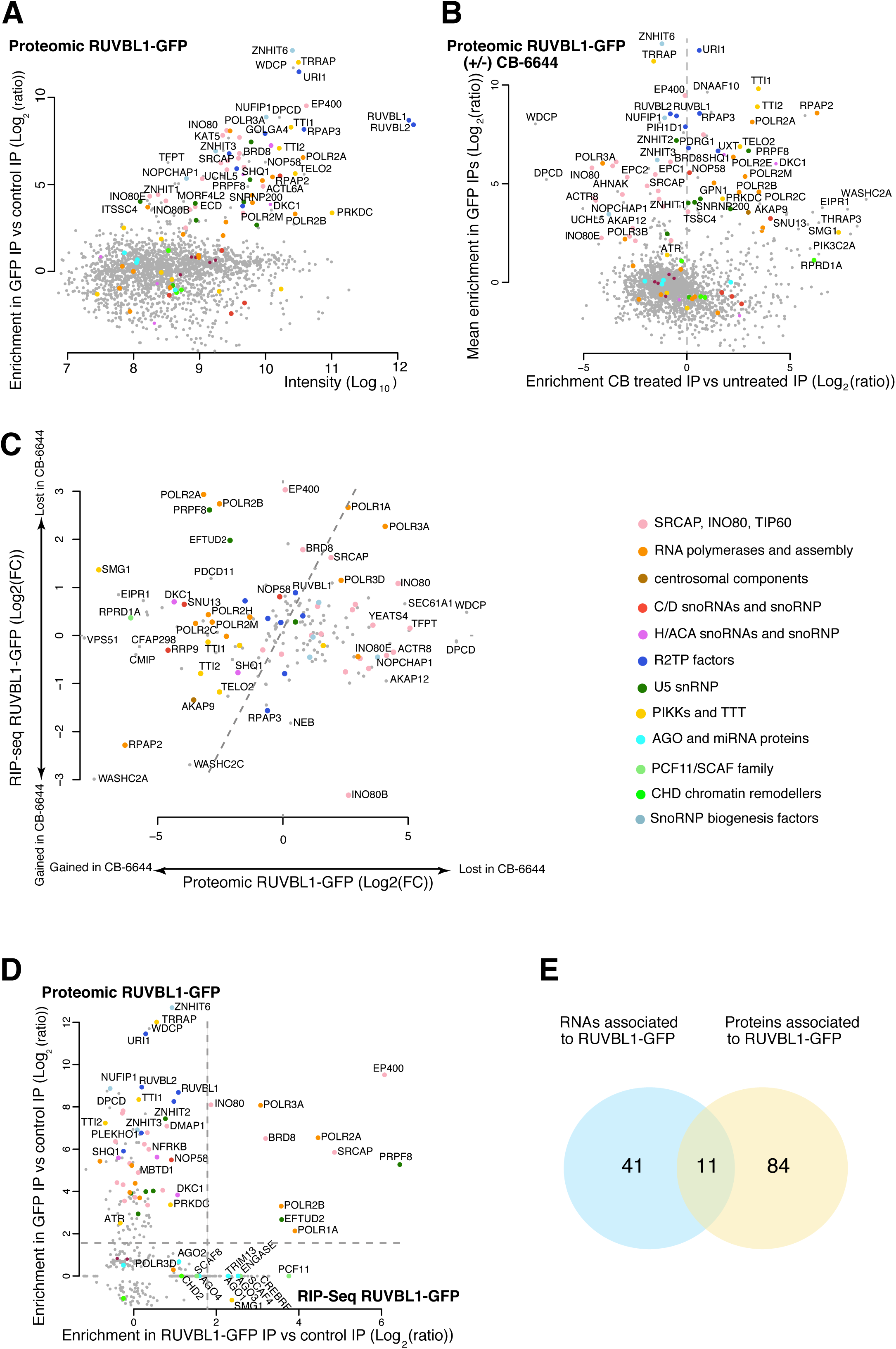
Comparison of the co- and post-translational interactomes of RUVBL1. **A-** Proteomic analysis of RUVBL1-GFP partners. The graph represents the enrichment in the GFP IP versus the control IP (y axis; Log2 (ratio)) as a function of signal intensity (x axis; Log10). Each dot represents a protein that is colored according to the code on the right of panel E. **B-** Proteomic comparison of RUVBL1-GFP partners in untreated and CB-6644 treated cells. The graph shows the mean enrichment in the two GFP IPs versus the control IPs (y axis, log2 (ratio)) as a function of the enrichment in the CB-6644 GFP IP over the untreated GFP IP (x axis, log2 (ratio)). **C-** Comparison of the effect of CB-6644 on the partners of RUVBL1-GFP found by RIP-seq and proteomics. The graph shows the fold change in the CB-6644 treated versus non-treated cells found in the RIP-seq (y axis, log2), as a function of the same fold change found by proteomic (y axis, log2). The fold change is computed as the (IP-GFP/IP-control) ratio in untreated cells, divided by the (IP-GFP/IP-control) ratio in CB-6644 treated cells. The top 100 RNA of the RIP-sq (best pValue) and the top 50 proteins (best pValue) are shown. **D-** Comparison of the RUVBL1-GFP partners found in the proteomic and RIP-seq experiment. The graph plots the enrichments in the RUVBL1-GFP IP versus the control IP, for the proteomics (y axis, log2 (ratio)), versus the RIP-Seq (x axis, log2 (ratio)). Only the top 200 partners with the best pValue in each IP are shown. **E-** Venn diagram. Blue and yellow discs are RUVBL1-GFP associated RNAs and proteins respectively. A partner is counted in the overlap if both its mRNA and protein have an enrichment more than 1.8 fold in the proteomic and the RIP-Seq experiments, respectively.

Interestingly, when we plotted the fold change induced by CB-6644 in the binding of RUVBL1 to mRNAs versus proteins (Figure 2C), it became apparent that the inhibition of RUVBL1/RUVBL2 ATPase activity had a differential effect, with many mRNAs losing binding upon CB-6644 treatment while the corresponding protein showed increased binding (POLR2A, POLR2B, PRPF8, EFTUD2, SMG1, DKC1). In addition, other subunits of these complexes also showed increased binding with CB-6644, as well as cofactors known to be involved in their assembly (POLR2C, POLR2H, POLR2M, RPAP2 for RNA polymerase II; TELO2, TTI1, TTI2, for SMG1; SHQ1 for DKC1). This indicated that RUVBL1/RUVBL2 followed an ATPase cycle linking co- and post-translation steps: they bind co-translationally these client mRNAs in an empty or ADP state, and require ATP hydrolysis to be released from later, post-translational assembly intermediates.

### RUVBL1 association to its partners can be strictly co-translational, strictly post-translational, or both co- and post-translational

Next, we wanted to determine which partners bind RUVBL1 co- or post-translationally, and we plotted the enrichments measured in the RIP-seq *versus* the AP-MS proteomic analysis, using the top 300 candidates in each of the two experiments made in untreated conditions (Figure 2D). This clearly identified three classes of RUVBL1 partners (Figure 2D-E): (i) only co-translational (41 partners), which spread along the x-axis and were enriched only in the RIP-seq experiments; (ii) only post-translational (84 partners), located along the y-axis and enriched only in the AP-MS proteomic analysis; (iii) co- and post-translational (11 partners), enriched in both experiments (Figure 2D). Interestingly, the latter corresponded to key subunits of the main R2TP client complexes (U5 snRNP, the RNA polymerases) as well as the chromatin remodeling complexes, while the post-translational partners corresponded mainly to the other subunits of these complexes and to R2TP cofactors. The strictly co-translational partners corresponded mostly to previously uncharacterized RUVBL1/RUVBL2 partners, indicating that the co-translational interactome of the R2TP chaperone is quite unique and not easily accessible by traditional proteomic approaches.

### RPAP3 binds clients co-translationally and associates with a similar set of polysomes as RUVBL1

RUVBL1/RUVBL2 are part of not only of R2TP but also several other complexes, such as the R2TP-like chaperones ^26^ and the INO80, SRCAP and TIP60 chromatin remodeling complexes. To determine whether the association of RUVBL1 to polysomes was taking place in the context of R2TP, we performed a RIP-seq experiment with RPAP3, another subunit of R2TP. We used CRISPR/Cas9 genome editing to generate a heterozygous HCT116 cell line with a GFP tag fused at the C-terminus of one allele of RPAP3. The cell line was characterized by PCR genotyping and Western blot, which confirmed the proper integration of the tag (Figure S2A-B). RNA sequencing of GFP IPs showed that 153 mRNAs were associated with RPAP3-GFP, as compared to a control IP (FDR<0.05; Table S1; Figure 3A). Comparison of the enrichment values measured in the RPAP3-GFP and RUVBL1-GFP IPs revealed a good correlation between the two experiments (Figure 3C): mRNAs highly enriched in the RUVBL1-GFP IP were also highly enriched in the RPAP3-GFP IP, with however some preferences. For instance, RUVBL1-GFP associated more strongly with C/D snoRNAs, SMG1 and EFTUD2 mRNAs, while RPAP3-GFP bound better the mRNAs coding for SNAPC4, the SCAF4/SCFA8/CHERP family of RNA polymerase II CTD binding proteins, and the CHD family of chromatin remodelers (Figure 3C). Interestingly, this was also the case for the PCNT and NIN mRNAs, which code for centrosomal proteins and which were recently shown to be locally translated at centrosomes, possibly to enable their co-translational assembly within centrosomes ^59^. Finally, although the presence of RUVBL1 in the INO80, SRCAP and TIP60 chromatin remodeling complexes is thought to be unrelated to R2TP ^60^, RPAP3-GFP was strongly associated with mRNAs coding key subunits of these complexes (EP400, SRCAP, INO80, BRD8). These mRNAs were the same that were bound by RUVBL1-GFP, raising the possibility that the R2TP chaperone could be involved in the biogenesis of these chromatin remodeling complexes, beyond the participation of RUVBL1/RUVBL2 in their mature forms.

**Figure 3.**
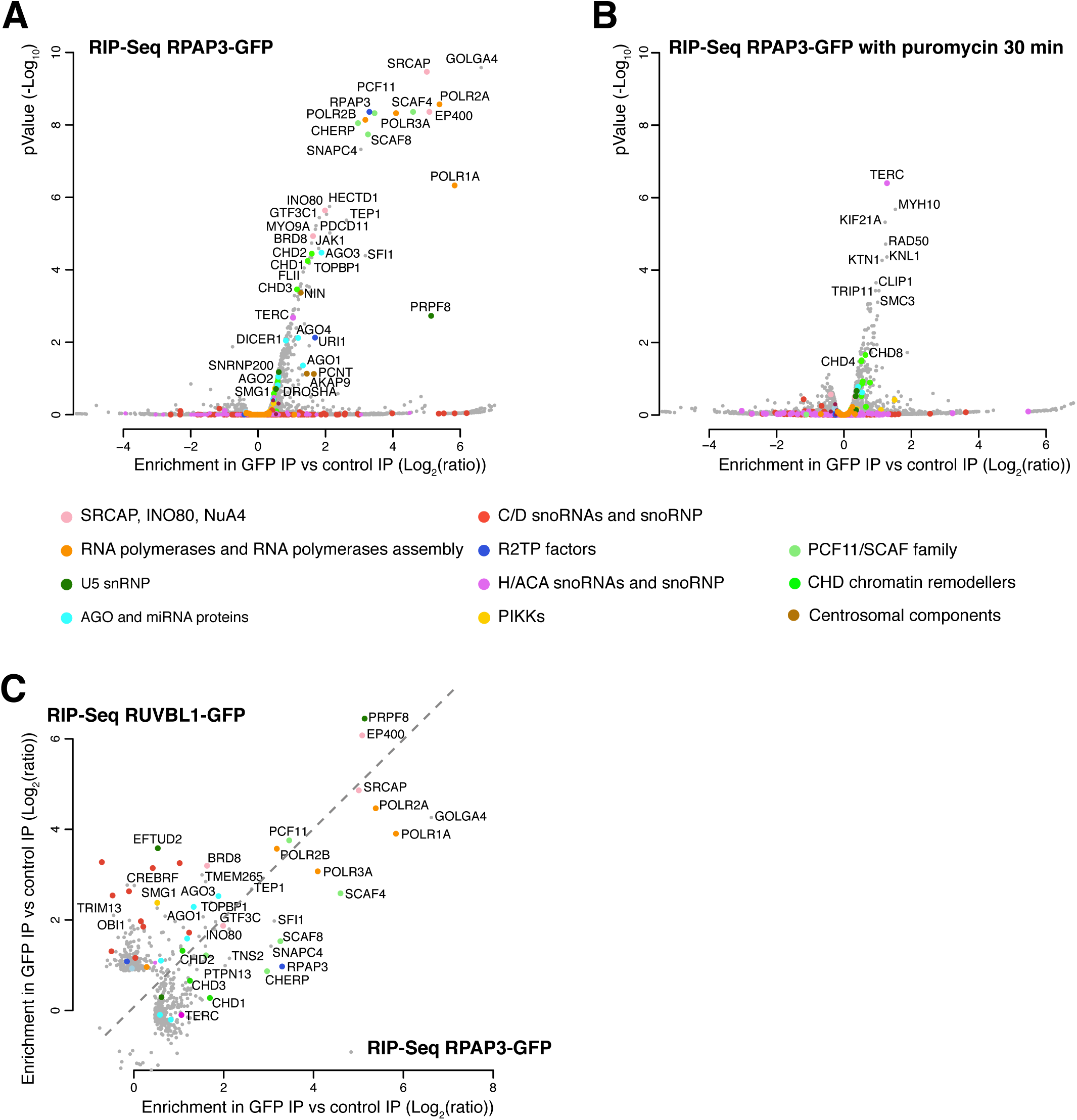
RPAP3 and RUVBL1 associate with similar RNAs. **A-** Volcano plot of the RPAP3-GFP RIP-seq, showing the pValue (y axis; -Log10) as a function of the enrichment in the GFP IP over the control IP (x axis; Log2(ratio)). Each dot represents a RNA that is colored according to the code below the panels A and B. **B-** Volcano plot of the RPAP3-GFP RIP-seq in cells treated with puromycin. Legend as in A. **C-** Comparison of the RUVBL1-GFP and RPAP3-GFP RIP-seq experiments. The graph displays the enrichment in the GFP IPs versus the control IPs, for the RUVBL1-GFP versus the RPAP3-GFP cells in untreated condition. Each dot is an RNA and is color coded as in A and B. Only the top 200 partners with the best pValue in each IP are shown.

Importantly, and as previously observed for RUVBL1, most mRNAs were lost or much less associated with RPAP3-GFP after a 30 min puromycin treatment (Figure 3B and S2C-D; Table S1). Only 20 out of the 153 enriched RNAs associated as well in puromycin as in untreated cells, and these generally had low enrichment values over the control IP (<1.5; Table S1). One interesting RNA of this class was TERC, the non-coding RNA of telomerase. This RNA carries an H/ACA motif and the R2TP chaperone is involved in telomerase assembly ^9,11,12,61^ suggesting that the translation independent RNA partners of RPAP3 could also be R2TP clients. In any case, these results demonstrated that the association of RUVBL1 with polysomes reflects an engagement of the full R2TP chaperone, and not a separate function of this protein.

### Artificial R2TP condensates recruit mRNAs coding for client proteins

It has been previously observed that interactions can form artifactually in extracts after cell lysis ^62^. To validate the interaction of R2TP with its target mRNAs, we thus used a co-recruitment assay to evaluate these interactions directly in intact cells ^13,16^. We generated artificial cytoplasmic condensates by using a fragment of DDX6 (amino-acids 289-483, or DDX6fm), sufficient to target the protein to P-bodies, and fusing it to DsRed2 and either RUVBL1 or RPAP3. RUVBL1 is normally diffusely present in the cytoplasm and the nucleoplasm (Figure 1B). In contrast, the DsRed2-DDX6fm-RUVBL1 fusion accumulated in cytoplasmic foci, creating a high local accumulation of R2TP. We then used smFISH to determine whether the mRNAs of interest were recruited to these foci (Figure 4). We tested some of the most-enriched mRNAs in the previous RIP-seq experiments, coding for: the polymerase subunits POLR2A, POLR2B, POLR1A, the U5 snRNP component PRPF8, and the subunits EP400 and BRD8 of the TIP60 histone acetylase complex. Remarkably, the endogenous POLR2A and PRPF8 mRNAs were efficiently recruited to the cytoplasmic foci formed by either DsRed2-DDX6fm-RUVBL1 or DsRed2-DDX6fm-RPAP3 (Figure 4A-B). In addition, this recruitment was lost after a 30 min puromycin treatment, and it did not occur with a DsRed2-DDX6fm-KPNA3, an unrelated protein used as a negative control (Figure 4C). To further confirm that the recruited mRNAs were translated, we performed a triple color experiment, adding an immuno-fluorescence labelling of the POLR2A protein (Figure 4D). Indeed, this protein also accumulated to the DsRed2-DDX6fm-RUVBL1 foci together with the POLR2A mRNA, and the protein label mostly disappeared after a 30 min incubation with puromycin. This confirmed that the recruited POLR2A mRNAs were translated. The other tested mRNAs were also recruited to DsRed2-DDX6fm-RUVBL1 cytoplasmic foci, although with a lower amount of mRNA per foci (Figure S3). Altogether, this validated the interactions observed in the RIP-seq experiments.

**Figure 4.**
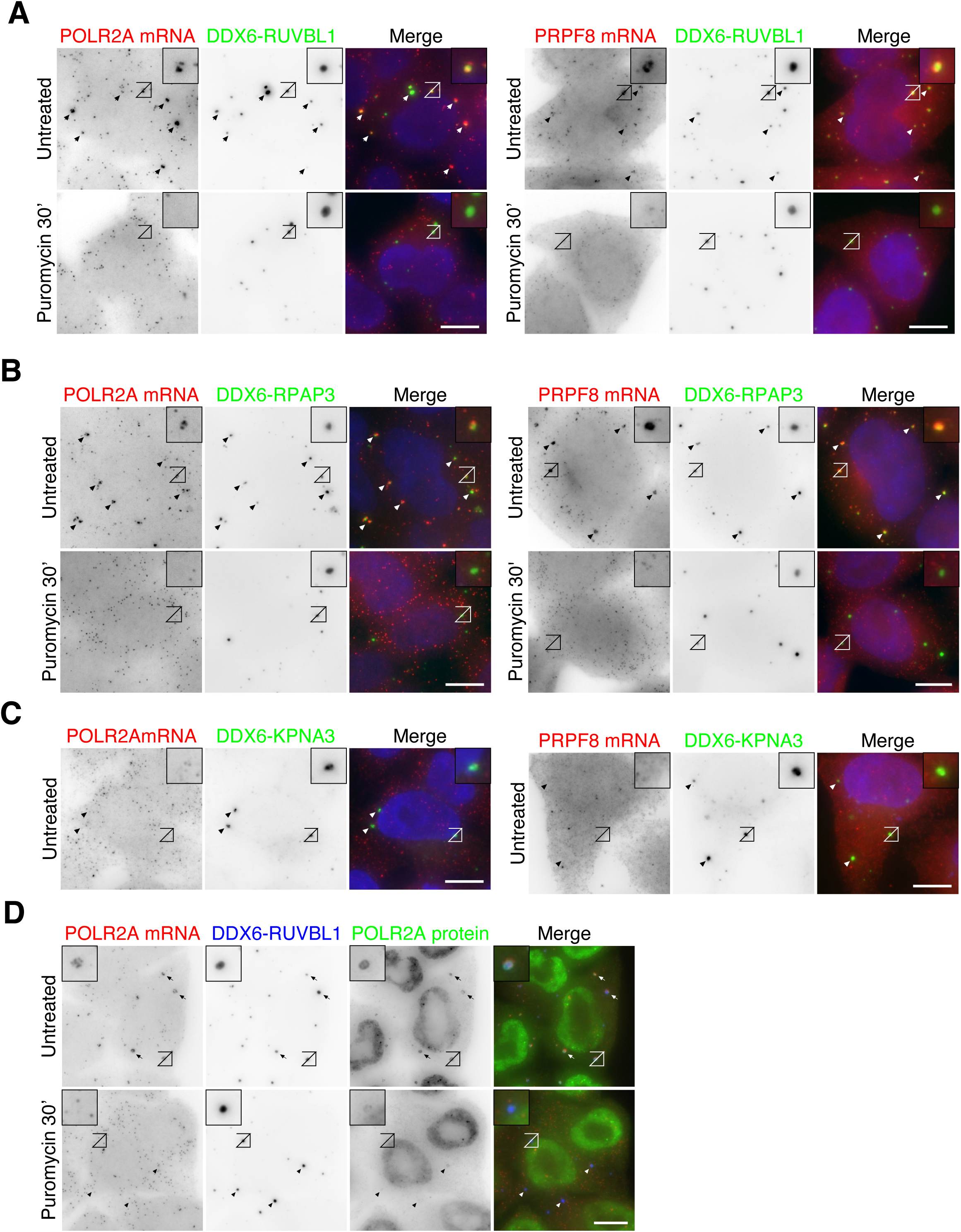
The POLR2A and PRPF8 mRNAs are recruited to artificial R2TP condensates. **A-** Micrographs of HeLa cells stably expressing DsRed2-DDX6fm-RUVBL1, treated with puromycin (bottom) or not (top), and labelled by smFISH for the POLR2A (left) or PRPF8 (right) mRNAs. Merge images (right of each panel) show mRNAs in red, DsRed2-DDX6fm-RUVBL1 in green and DAPI staining in blue. Scale bar: 10 µm. Arrowheads point to the location of artificial R2TP condensates in the cytoplasm. Insets show a higher magnification of the boxed area. **B-** Legend as in A but with DsRed2-DDX6fm-RPAP3. **C-** Legend as in A but with the negative control with DsRed2-DDX6fm-KPNA3. **D-** Micrograph of HeLa cells stably expressing DsRed2-DDX6fm-RUVBL1 treated or not with puromycin and labelled for POLR2A mRNAs by smFISH (left) and the endogenous POLR2A protein by immunofluorescence (middle right). Merge images show POLR2A mRNAs in red, DsRed2-DDX6fm-RUVBL1 in blue and the POLR2A protein in green. Scale bar: 10 µm. Arrowheads point to artificial R2TP condensates in the cytoplasm. Inset shows a higher magnification of the boxed area.

### Single molecule imaging reveals R2TP factories accumulating POLR2A and PRPF8 mRNAs

Next, we wanted to quantify the interactions of R2TP with mRNAs in a more native context, and we thus tagged the endogenous RPAP3 with the SunTag. Indeed, this is a repeated epitope that enables single protein detection in native cells with a recombinant mono-chain antibody fused to GFP ^63^. Genotyping and Western blots confirmed the proper heterozygous integration of the SunTag in HeLa cells, and revealed that the RPAP3-SunTag was much less expressed than its untagged counterpart (∼50 fold less, Figure S4). Interestingly, fluorescence microscopy showed that the RPAP3-SunTag could be observed not only as single proteins in the cytoplasm but also as foci containing multiple molecules (Figure 5A). We then performed smFISH against POLR2A and PRPF8 mRNAs. Surprisingly, both of these mRNAs were present as single molecules and also as foci, which colocalized with the RPAP3-SunTag foci (Figure 5A). Counting both the mRNAs present as foci and single molecules, we measured that an average of 30% and 35% of the PRPF8 and POLR2A mRNAs colocalized with RPAP3-SunTag, respectively (Figure 5B). After treatment with puromycin for 30 minutes, the foci containing RPAP3-SunTag, POLR2A and PRPF8 mRNA disappeared and the colocalization of these mRNAs with RPAP3-SunTag dropped significantly (Figure 5A-B). Importantly, the colocalization of POLR2A and PRPF8 mRNAs with RPAP3-SunTag occurred not only in the foci but also when these mRNAs were present as single molecules (Figure S4C). This colocalization appeared lower than in the foci (20% versus 75%; Figure S4C and 4D), but it was still significant as it decreased after puromycin treatment. The relatively high remaining colocalization in puromycin-treated cells (16% for POLR2A mRNA, Figure S4C) likely reflected background events due to the density of the RPAP3-SunTag labeling. Indeed, similar values were obtained with alpha-catenin mRNA, a control unrelated to R2TP. The other mRNAs did not form foci (POLR2B, EFTUD2, BRD8, EP400). Nevertheless, we could detect a puromycin-dependent colocalization with RPAP3-SunTag for all of them, which was not the case for the control alpha-catenin mRNA (Figure 5B). Their colocalization was lower than for POLR2A and PRPF8 mRNAs (15-25% vs. 30-35%), but the large excess of untagged RPAP3 (50 fold) is likely to underestimate the true colocalization values.

**Figure 5.**
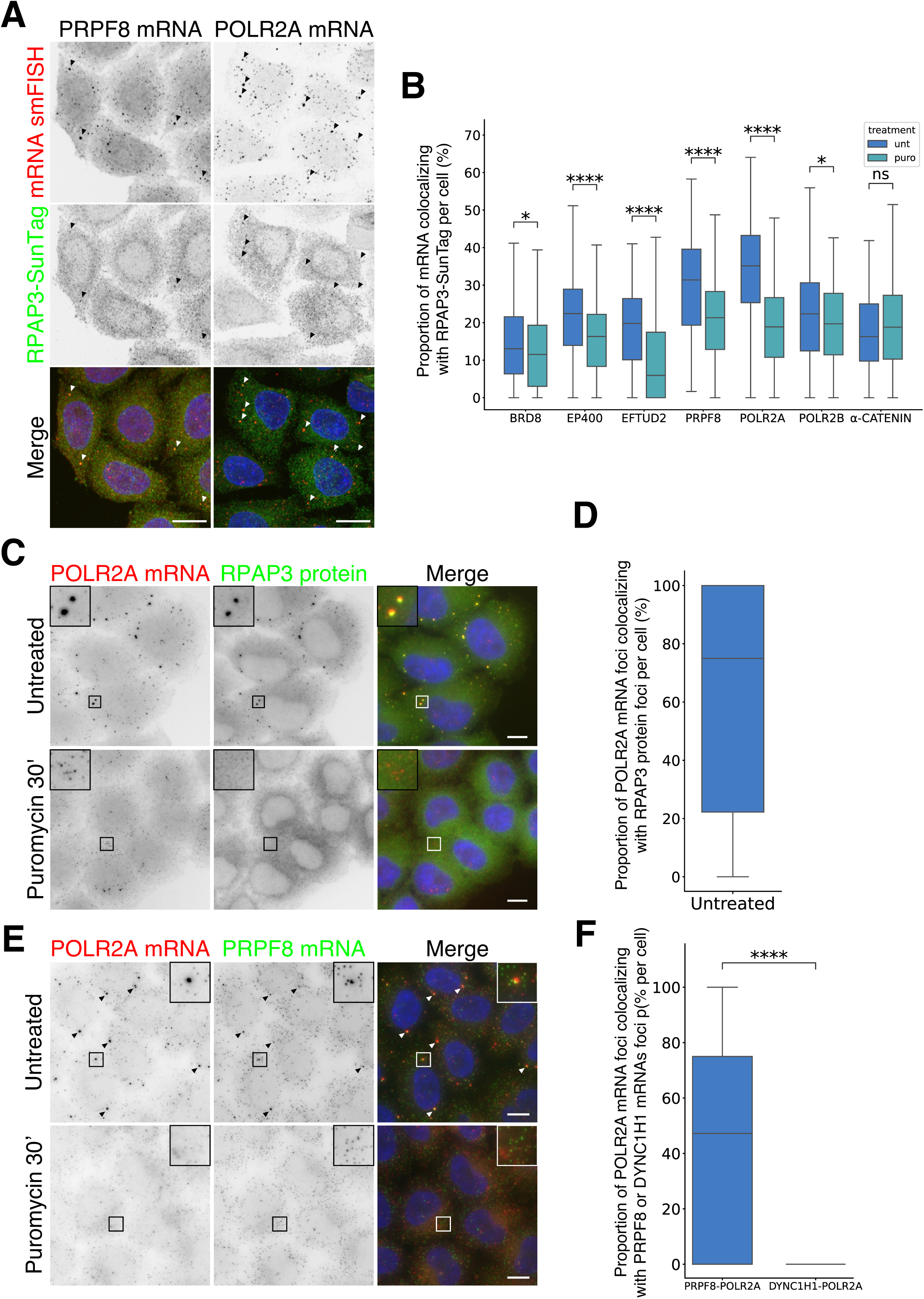
Colocalization of RPAP3 protein, PRPF8 and POLR2A mRNA in cytoplasmic foci. **A-** Micrographs of HeLa Flp-In cells expressing a RPAP3-32xSuntag allele (middle) and labelled for POLR2A or PRPF8 mRNAs by smFISH (top). Merge images (bottom) show mRNAs in red, RPAP3-SunTag in green and DAPI staining in blue. Scales bars are 20µm. **B-** Quantification of the colocalization of RPAP3-SunTag with the indicated mRNAs. Boxplots represent the proportion (y axis, %) of several mRNA detected by smFISH and colocalizing with RPAP3-SunTag under normal (unt, blue) and puromycin conditions (puro, light blue). Each observation is a cell, at least 200 cells were analyzed per measurements. Within each box, the middle horizontal line shows median value; boxes extend from the 25th to the 75th percentile of each group’s distribution of values; vertical extending lines lowest and highest 25th percentile of the distribution. P-values were determined using Welch’s t-test. * P<0.05, ** P<0.01, *** P<0.001, *** P<0.0001. **C-** Micrographs of HeLa Kyoto cells treated or not with puromycin and labelled for the POLR2A mRNA by smFISH (left) and for the endogenous RPAP3 protein by immunofluorescence (middle). Merge images (right) show POLR2A mRNA in red, RPAP3 protein in green and DAPI staining in blue. Scale bars are 10µm. Inset shows a higher magnification of the boxed area. **D-** Quantification of the colocalization of POLR2A mRNA foci with the endogenous RPAP3 protein foci. Boxplots represent the proportion (y axis, %) of POLR2A mRNA colocalizing with RPAP3 foci under normal conditions. Each observation is a cell, 86 cells were analyzed. **E-** Micrographs of HeLa Kyoto cells treated or not with puromycin, and labelled for POLR2A (left) and PRPF8 (middle) mRNAs by smFISH. Merge images (right) show POLR2A mRNA in red, PRPF8 mRNA in green and DAPI staining in blue. Scale bars are 10µm. Inset shows a higher magnification of the boxed area. **F-** Quantification of the proportion of POLR2A mRNA foci colocalizing with either DYNC1H1 mRNA foci or PRPF8 mRNA foci in HeLa Kyoto cells, treated or not with puromycin. Legend as in D. Each observation is a cell, at least 125 cells were analyzed per condition.

The foci observed with either RPAP3-SunTag or the POLR2A and PRPF8 mRNAs were intriguing as they suggested the existence of structures dedicated to the translation and co-translational binding of these proteins to the R2TP chaperone. To explore this possibility, we used native, untagged HeLa Kyoto cells and labelled endogenous RPAP3 by immunofluorescence together with POLR2A mRNAs. Remarkably, both the POLR2A mRNA and RPAP3 formed foci and these foci co-localized (Figure 5C-D). On average, cells contained four RPAP3 foci and 75% of the POLR2A foci colocalized with RPAP3 (Figure 5D and S4D-E). Moreover, treatment with puromycin for 30 minutes led to the near total disappearance of either type of foci, indicating that their formation is driven by the nascent protein chains (Figure S4D-E). We then tested PRPF8 mRNA, which also formed foci in the RPAP3-SunTag cells. Indeed, this mRNA formed foci in native, untagged HeLa Kyoto cells, which were also puromycin sensitive and which frequently co-localized with POLR2A mRNAs foci (∼50%, Figure 5E-F and S4D-E). Several mRNAs were recently reported to form translation factories, which are small cytoplasmic foci accumulating specific mRNAs undergoing translation and sensitive to puromycin inhibition ^64^. Two color smFISH showed that the POLR2A mRNA foci did not colocalize with the DYNC1H1 translation factories, which is not related to the R2TP (Figure 5F and S4F). However, the POLR2A mRNA foci stained for the POLR2A protein and the staining disappeared with puromycin (Figure S4G and H), indicating that the POLR2A foci are indeed translation sites. Because they also accumulate RPAP3, these foci thus appeared to be specialized for both the translation of specific mRNAs and R2TP activity.

### RNAs coding for pairs of subunits that assemble together rarely colocalize

Recent evidence suggests that mRNA coding for proteins that assemble together can in some cases be colocalized and co-translated, a pathway termed co-co assembly ^45,52^. The colocalization of POLR2A and PRPF8 mRNAs in foci thus prompted us to test additional mRNA pairs, as our R2TP RIP-Seq experiments revealed several mRNAs coding for pairs of subunits belonging to the same complex, such as POLR2A/POLR2B for RNA polymerase II, PRPF8/EFTUD2 for U5 snRNP and EP400/BDR8 for the TIP60 chromatin remodeling complex (Figure 6A). Surprisingly, dual-color smFISH showed only a very limited colocalization between the mRNA pairs (Figure 6B-C). The highest level was observed for BRD8 and EP400, with an average of 5% of BRD8 mRNA colocalizing with EP400 mRNA. This low level nevertheless appeared significant because it dropped to less than 1% after a 30 min puromycin treatment, similar to a control JUP mRNA. We also observed a low level of colocalization, but still significant, between POLR2B and POLR2A mRNAs (3.5%; Figure 6B and C), while no significant colocalization was observed between EFTUD2 and PRPF8 mRNAs. Altogether, these data are not in favor of the co-co assembly model where polysomes of subunit pairs associate together. This was further reinforced by the fact that the POLR2B and POLR1A mRNAs colocalized to the same extent with POLR2A mRNAs, even though they belong to different protein complexes (Figure S5). This suggested that the low level of mRNA colocalization that we observed is not related to co-co assembly.

**Figure 6.**
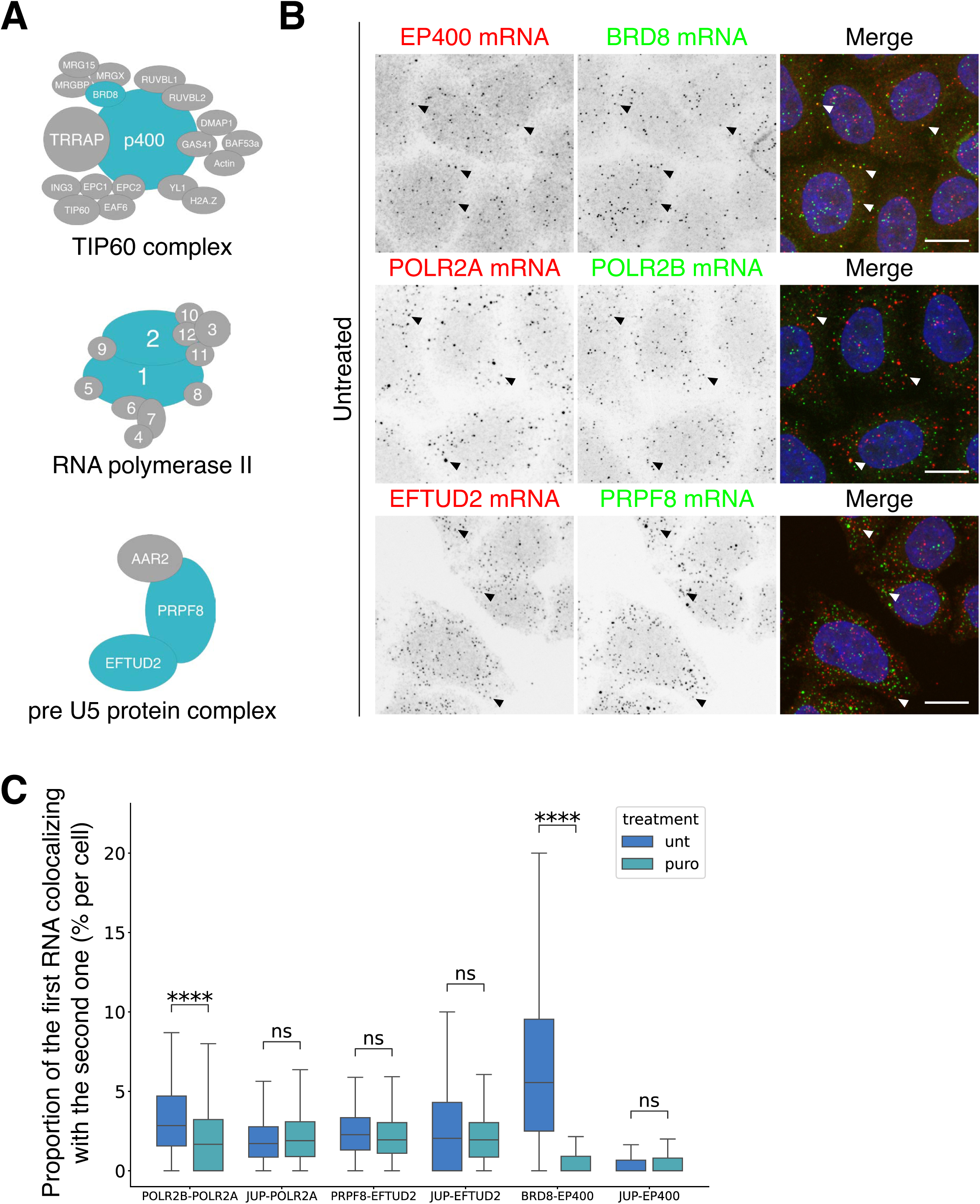
Pairs of mRNAs coding for proteins that assemble together show little colocalization. **A-** Schematics of the protein complexes. **B-** Micrographs of HeLa Flp-In cells labelled by dual color smFISH for the indicated mRNAs (left: EP400, POLR2A, EFTUD2, middle: BRD8, POLR2B, PRPF8). Merge images (right) show left mRNAs in red, right mRNAs in green and DAPI staining in blue. Arrowheads point to cytoplasmic mRNA foci. Scale bars are 20µm. **C-** Quantification of the colocalization for the pairs of mRNAs shown in B and with JUP as a negative control mRNA. Boxplots represent the proportion of mRNA colocalized (y axis, %) s under normal (unt, blue) and puromcyin conditions (puro, light blue). Each observation is a cell, at least 200 cells were analyzed except for JUP-EFTUD2 (32 cells). Within each box, the middle horizontal line shows the median value; the box extends from the 25th to the 75th percentile of the distribution of values; vertical lines extend to the lowest and highest percentile of the distribution. P-values were determined using Welch’s t-test. * P<0.05, ** P<0.01, *** P<0.001, *** P<0.0001.

### The HSP90 and RUVBL1/2 ATPase activity are required for localized mRNA translation

The accumulation of the RPAP3 protein in cytoplasmic foci, together with POLR2A and PRPF8 mRNAs, suggests that these structures function as R2TP factories to enhance interaction of nascent chains with the chaperone via a proximity effect. The formation of these structures depend on translation (Figure S4D-E) and we wondered whether the chaperones, which interact with the nascent proteins, would themselves be involved in their formation. To address this question, we inhibited RUVBL1/RUVBL2 ATPase activity with CB-6644 and analyzed POLR2A mRNA localization (Figure 7A and B). Interestingly, this led to a diminution of the number of POLR2A mRNA foci after 4h and 8h of treatment, and their complete disappearance after 16h. The same effect was observed for PRPF8 mRNAs and remarkably, the mRNA foci also disappeared when the ATPase activity of HSP90 was inhibited with Geldanamycin for 1h (Figure 7C-E). Thus, the HSP90 and RUVBL1/RUVBL2 chaperones play an essential role in the formation of the POLR2A and PRPF8 translation factories, indicating that they not only interact with the client proteins but also control their localized translation.

**Figure 7.**
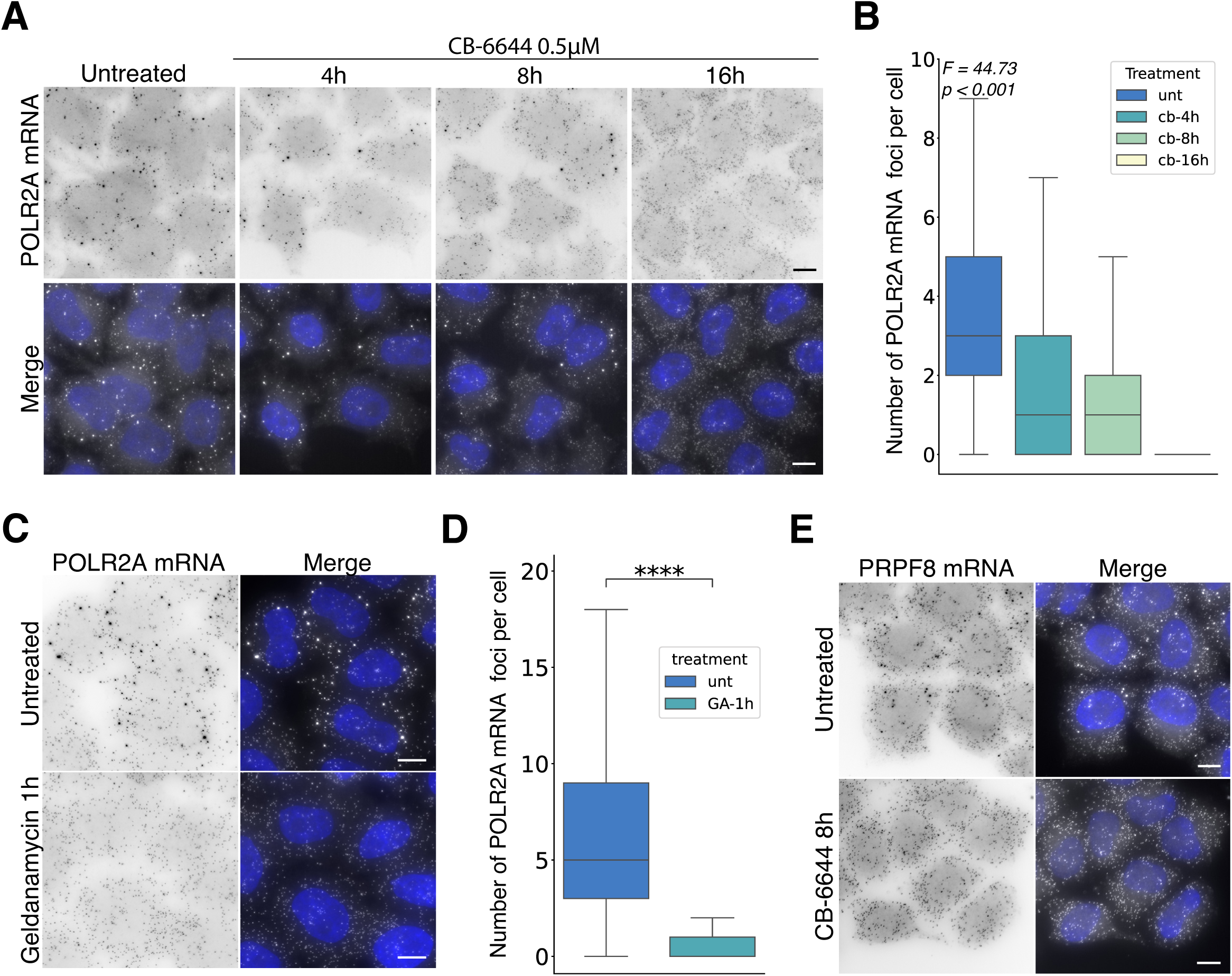
Inhibition of HSP90 and RUVBL1/2 ATPase activity leads to the disassembly of POLR2A and PRPF8 mRNA foci. **A-** Micrographs of HeLa Kyoto cells labelled by smFISH for POLR2A mRNA (top) in cells untreated or treated with CB-6644 for 4h, 8h and 16h. Merge images (bottom) show mRNAs in white and DAPI staining in blue. Scale bars are 10µm. **B-** Quantification of the number of POLR2A mRNA cytoplasmic foci in cells upon CB-6644 treatment. Boxplot represents the number of POLR2A mRNA cytoplasmic foci (y axis) under normal (unt, blue) and CB-6644 (CB-4h, light blue; CB-8h, light green; CB-16h, light yellow) conditions. Each observation is a cell, at least 110 cells were analyzed. Variability among all the groups was determined by performing one-way ANOVA (displayed statistics and p-value). **C-** Micrographs of HeLa Kyoto cells labelled by smFISH for POLR2A mRNAs (left) in cells untreated or treated with Geldanamycin for 1h. Merge images (right) show mRNA in white and DAPI staining in blue. Scale bars are 10µm. **D-** Quantification of the number of POLR2A mRNA foci per cell upon Geldanamycin treatment. Boxplot represents the number of POLR2A mRNA foci per cell (y axis) under normal (unt, blue) and Geldanamycin (GA-1h, light blue) conditions. Each observation is a cell, at least 115 cells were analyzed. P-values were determined using Welch’s t-test. * P<0.05, ** P<0.01, *** P<0.001, *** P<0.0001. **E-** Micrographs of HeLa Kyoto cells labelled by smFISH for PRPF8 mRNAs in cells untreated (top) or treated with CB-6644 (bottom) cells for 8h. Scale bars are 10µm. Merge images (right) show mRNAs in white and DAPI staining in blue.

Interestingly, the previous RIP-Seq experiments raised the possibility that the R2TP chaperone could be involved in another case of local translation, as the RPAP3-GFP pulled down several mRNAs localized near the centrosomes: NIN, PCNT and AKAP9 (see Figure 3). Moreover, other centrosomal mRNAs were among the top hits in the RPAP3 RIP-seq (Table S1), although their enrichment was not individually statistically significant: CEP350 (ranked 321 out of ∼18000; Table S1), CCDC88C (534), BICD2 (1111), NUMA1 (1415) and ASPM (1673). In fact, out of the 8 mRNAs known to localize to human centrosomes in a translation-dependent manner ^59^, all but one were found in the top hits, representing a very significant enrichment for the entire mRNA family (pValue < 0.001). To test whether the R2TP plays a role in the local translation of centrosomal mRNAs, we measured by smFISH the centrosomal localization of NIN mRNAs with and without treatment with CB-6644 (Figure 8A-B). We found a nearly 2-fold decrease in the amount of NIN mRNAs at centrosomes upon treatment, indicating a role of the RUVBL1/RUVBL2 ATPases in its localized translation. Altogether, these data demonstrated an important role of the R2TP chaperone in the control of the local translation of its mRNA targets.

**Figure 8.**
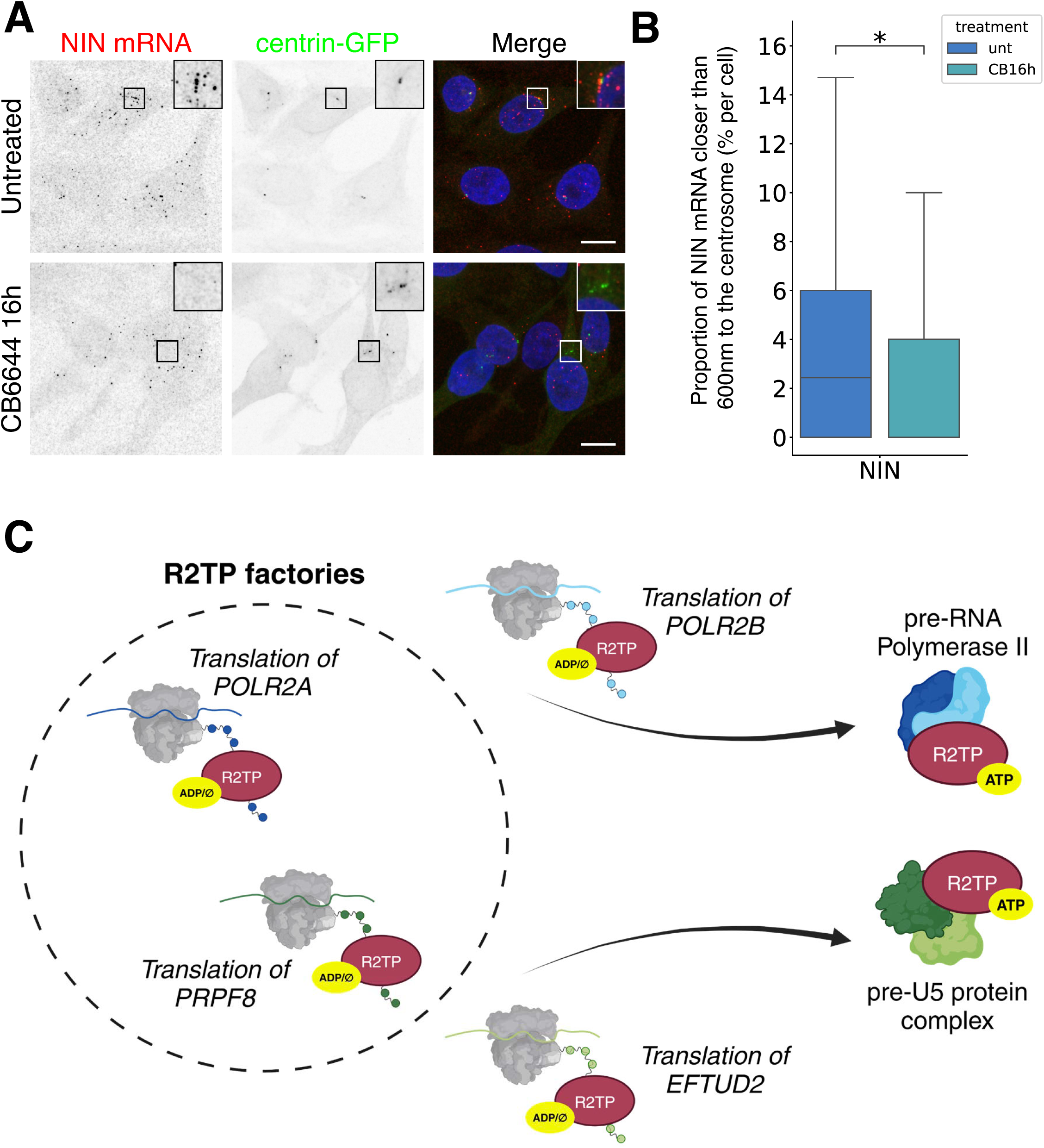
Inhibition of the R2TP ATPase activity inhibits the centrosomal localization of NIN mRNAs. **A-** Micrographs of HeLa cells stably expressing centring-GFP (middle), treated or not with CB-6644 and labelled for NIN mRNAs by smFISH (left). Merge images (right) show mRNAs in red, centrin-GFP in green and DAPI staining in blue. Scale bars are 20µm. **B-** Quantification of the localization of NIN mRNAs to the centrosome. Boxplots represent the proportion of mRNA closer than 600nm to the centrosome (y axis, %) under normal (unt, blue) and CB-6644 (CB-16h, light blue) conditions. Each observation is a cell, at least 210 cells were analyzed. P-values were determined using Welch’s t-test. * P<0.05, ** P<0.01, *** P<0.001, *** P<0.0001. **C-** Model describing the cha-cha assembly pathway and the R2TP cycle between co- and post-translational steps (see text).

## Discussion

Assembly of protein complexes has recently been shown to involve co-translational steps ^47^. One free, fully translated subunit may bind another subunit while it is translated (co-post pathway), or two mRNAs coding for partner subunits may associate to be co-translated, and the subunits co-assembled during their translation (co-co pathway). These questions of co-translational assembly have generally not been investigated in the context of chaperone-mediated assembly. Here, we studied the HSP90/R2TP chaperone and surprisingly, we found little evidence for the co-co pathway. Instead, the fate of client subunits appears to be determined co-translationally by the loading of the R2TP assembly chaperone in dedicated factories.

### The R2TP assembly chaperone binds many of its client in a co-translational manner

Our RIP-seq experiment using RUVBL1-GFP and RPAP3-GFP knock-in cell lines shows that the HSP90/R2TP chaperone binds its client while they are being translated. We found that ∼150 mRNAs are bound by R2TP in a translation-dependent manner but this is likely an underestimate because some mRNAs coding for known R2TP clients are in the top ranks of the IPs although they do not pass the statistical threshold. Overall, this indicates that the co-translational binding of R2TP to its clients is widespread. This includes well known clients such as subunits of the RNA polymerases or the U5 snRNP and factors previously known to associate with RUVBL1/RUVBL2 but not as R2TP clients, such as subunits of the chromatin remodeling complexes INO80, SRCAP and TIP60. The co-translational interactions of R2TP also reveal many novel partners and this is quite remarkable as RUVBL1/RUVBL2 have been heavily studied, including by a number of large-scale interactomic approaches ^21,22,23,24^. This indicates that co-translational interactions represent a part of the interactome that is difficult to access by more traditional methods.

It is noteworthy that RUVBL1/RUVBL2 not only associate with polysomes but also cross-link with polyA+ mRNAs ^65^, raising the possibility they may be atypical RNA binding proteins. However, the fact that translation inhibition disrupts most of their interactions with mRNAs suggests that their crosslinks with polyA+ RNA may be due to a proximity effect upon nascent protein binding, rather than a traditional sequence-specific interaction. Interestingly, some RNAs remained associated with R2TP in absence of translation. This was the case for a few mRNAs, although their binding was generally weak, and for many non-coding RNAs such as the C/D snoRNAs and the telomerase RNA. Remarkably, inhibition of the ATPase activity of RUVBL1/RUVBL2 strengthened their binding to C/D snoRNAs and to the core C/D protein SNU13, indicating that ATP hydrolysis is required for their release from immature C/D snoRNPs. Their dissociation could trigger the release of additional assembly factors, similar to the role of R2TP in releasing SHQ1 from DKC1 ^11^, and this could drive the final maturation of C/D snoRNPs.

In contrast to C/D snoRNAs, inhibition of the ATPase activity of RUVBL1/RUVBL2 dissociated them from most mRNAs, with some exceptions such as the WASHC2A/WASHC2C mRNAs and the PCF11/SCAF family. The fact that RUVBL1 associates much less with a series of client mRNAs when ATP-locked, while at the same time binding more strongly to the corresponding proteins (POLR2A, POLR2B, PRPF8, SMG1, DKC1, NOP58; Figure 2E), suggests a cycle in which RUVBL1/RUVBL2 are loaded co-translationally on their client in an apo state and remain associated post-translationally in an ATP state, probably as part of a partially assembled intermediate containing additional subunits and cofactors. Possibly, ATP hydrolysis may be required here to remodel these assembly intermediates and release RUVBL1/RUVBL2 and the other assembly factors. The ATP cycle of RUVBL1/RUVBL2 thus regulates co-translational client loading and release following completion of assembly, thereby coordinating co- and post-translational steps.

### The co-translational and post-translational interactomes of R2TP are distinct

A comparison of the RIP-Seq of RUVBL1-GFP with the corresponding proteomic data identifies three classes of partners, which bind RUVBL1: (i) only co-translationally, (ii) only post-translationally, or (iii) both co- and post-translationally. Only a minority of partners is bound both co- and post-translationally, indicating that the co-translational interactome of R2TP is distinct and quite specific, thus revealing a new facet of this important chaperone. Post-translational partners include proteins that bind indirectly to RUVBL1, such as some subunits of the RNA polymerases or the chromatin remodeling complexes, and cofactors involved in assisting assembly of key client complexes, such as ECD/TSSC4 for U5 snRNP, SHQ1 for H/ACA snoRNPs and ZNHIT3/ZNHIT6/NUFIP1/NOPCHAP1 for C/D snoRNPs. Possibly, some of these assisting factors may themselves bind co-translationally to their client proteins, chaperoning them and facilitating R2TP loading as shown for NOPCHAP1 ^42^. Proteins that require to be fully folded to associate with RUVBL1 should also fall in this category. This may be the case for PIKKs because while the R2TP is involved in their biogenesis ^18,19^, we did not observe co-translational interactions of RPAP3 or RUVBL1 with their mRNAs, with the exception of SMG1. The PIKKs have been shown in yeast to recruit the TTT complex co-translationally, to enable folding of the C-terminal hydrophobic amino-acids back into the protein core ^66^. The TTT recruits the R2TP via a phospho-dependent interaction of TELO2 with PIH1D1 ^41^, but this may occur post-translationally after the complete folding of the PIKKs, as suggested for their other partners ^66^.

Proteins bound both co- and post-translationally by R2TP include key subunits of its best characterized client complexes, such as the largest subunits of the nuclear RNA polymerases and key components of the U5 snRNP. Although the INO80, SRCAP and TIP60 chromatin remodeling complexes are not usually classified as R2TP clients, some of their key subunits of also fall in this category. All these proteins are large subunits of nuclear complexes that, for U5 snRNP and RNA polymerase II, require many assembly factors ^60^. These subunits are thus likely accompanied by the chaperone during a large part of their biogenesis and in agreement, chaperone deficiencies induce their retention in the cytoplasm ^16,67^. Surprisingly, we found many R2TP partners that are only detected in the co-translational dataset. They often constitute protein families, as for instance the AGO proteins, the PCF11/SCAF family and the CHD chromatin remodelers. This suggests a very early chaperone function of R2TP for these clients, perhaps to deposit a partner co-translationally or to help folding the protein. In this regard, it is interesting to note that the AGO proteins have hydrophobic C-terminal residues that fold back in the protein core much like the PIKKs ^68^. As previously proposed ^66^, co-translational chaperone binding may be especially important to maintain these proteins in a folding competent state until the completion of translation.

### The HS90/R2TP chaperone assembles translation factories and regulates local translation

The assembly of proteins may be favored by the physical association of their polysomes ^47^ and the so-called co-co pathway has recently been proposed to be widespread ^52^. The R2TP is well known to function in the assembly of protein complexes and we found here that it associates co-translationally with pairs of subunits that assemble together, such as POLR2A/POLR2B, PRPF8/EFTUD2 and EP400/BRD8. Surprisingly, we found only little colocalization of these mRNAs, indicating that the co-co pathway is marginal. Instead, we found that POLR2A and PRPF8 mRNAs concentrate in cytoplasmic foci that also accumulate R2TP, which we thus termed R2TP factories. These foci require ongoing translation and remarkably, also the HSP90 and R2TP ATPase activity. Polysomes have multiple nascent protein chains and RUVBL1/RUVBL2 form heteromeric hexamers, thus enabling multivalent interactions that could drive condensate formation. Moreover, RPAP3 can bind multiple times to RUVBL1/RUVBL2 and its N-terminal domain can also multimerize ^35^, enhancing the valency of the system. Because HSP90 directly binds RPAP3 and client proteins, the multivalency of R2TP provides a possible explanation for the role of HSP90 in the condensation of these polysomes.

The mRNA foci induced by HSP90 and R2TP are heterotypic in the sense that they can contain both POLR2A and PRPF8 polysomes, but they lack other mRNAs that also associate with R2TP. It may be that POLR2A and PRPF8 have particularities favoring polysome condensation, such as a stronger or higher multivalent binding of the nascent proteins to HSP90 and R2TP, or a controlled ribosomal pausing providing more time for binding the chaperones and condensing the polysomes. In agreement with this idea, it has been observed that unfolded protein domains can induce ribosome pausing and that this can sometimes drive polysome condensation ^69^. It is for instance the case for the CNOT1 granules that form under stress and accumulate the mRNAs coding for Rpt1 and Rpt2 proteasomal subunits, which then co-assemble ^69^. Importantly, the R2TP factories reported here are present in native cells and they do not contain mRNA pairs, ruling out that they function in the co-co assembly pathway. Rather, they appear to channel assembly by concentrating the nascent protein chains and chaperones, promoting co-translational interactions. This provides an alternative model of assembly, which we termed channeling by chaperone (cha-cha). In this model, the joining of subunits is determined co-translationally by the loading of the appropriate assembly chaperones, enabling assembly itself to occur elsewhere (Figure 8C).

## Material and methods

### Cell culture and drug treatments

HCT116 cells were grown in McCoy’s 5A medium containing 10% fetal bovine serum (FBS, Sigma-Aldrich) and 1% penicillin/streptomycin (Thermofisher) at 37°C, 5% CO_2_. HeLa Flp-In and HeLa Kyoto cells were grown in DMEM medium instead of McCoy’s 5A. All cells were routinely tested for the absence of mycoplasma. For cell treatments, drugs were used as follow: puromycin at 100 µg/mL for 30 min, CB-6644 at 500 nM for 4 to 16 h, Geldanamycin at 2 µM for 1h.

### Plasmids and transfection for co-recruitment assays

Plasmids were constructed using the Gateway technology (Invitrogen) with donor vectors from the human ORFeome ^70^. Detailed maps are available upon request. Stable cell lines expressing DsRed2-DDX6fm-RUVBL1 or DsRed2-DDX6fm-RPAP3 were obtained using the Flp-In system (Thermo-Fischer Scientific). Transient transfection was performed for the DsRed2-DDX6fm-KPNA3 construct. Vectors were transfected using JetPrime reagent and buffer (Polyplus) according to the manufacturer instructions: 200µL JetPrime Buffer, 2µg vector DNA and 4µL Jetprime reagent for cells cultured in 2mL in 6-well plates.

### Insertion of GFP and SunTag cassettes by CRISPR/Cas9 knock-in

HCT116 cells were CRISPR/Cas9 edited to insert a mAID-GFP-IRES-Neomycin cassette in fusion to the C-termini of RPAP3 and RUVBL1. The recombination cassettes contained 500 bases of homology arms flanking a cassette composed of mAID-GFP, a stop codon, an IRES containing a start codon fused to a Neomycin resistance gene and a stop codon. HeLa Flp-In cells stably expressing the SunTag scFv-sfGFP were CRISPR/Cas9 edited to insert a SunTag*32 cassette at the C-terminus of RPAP3. Cells were transfected using JetPrime (Polyplus) as above, with a cocktail of four plasmids including the repair cassette and constructs expressing the Cas9-nickase and two guide RNAs. Cells were selected on 800 µg/ml G418 neomycin for a few weeks. Individual clones were then picked and analyzed for GFP expression by fluorescence microscopy, Western blotting and PCR genotyping. The sequences targeted by the guide RNAs were (PAM sequences are lowercase): TCAGCAGTAAGAGACTCCCCagg for RUVBL1 guide 1, ACTTCATGTACTTATCCTGCtgg for RUVBL1 guide 2; GAGCTGAAAAAGAGGTATGGtgg for the RPAP3 guide. PCR genotyping was conducted using Phusion polymerase (NewEngland Biolabs) and genomic DNA prepared with Wizard® Genomic DNA Purification Kit (Promega). The sequences of the oligonucleotides were as follows (5’ to 3’). Fw1 in the gene before the homology arms: GGTGAAGTCCCCATGTCCTG for RUVBL1 and TGTCGACCGATCAGTGGTAGTAA for RPAP3; Rev1 in the mAID-GFP cassette: CGGTGCTCCGTCCATTGATA; Rev2 in the SunTag*32 cassette: ACTAGTGACCGGTCCTCTAGA; Fw2 in the repair cassettes: GTTCATAAACGCGGGGTTCG; Rev2 in the gene after the homology arm: CGCAGCTCTCATCCTACTGG for RUVBL1 and ACGCTACACTCACATCCAACA for RPAP3.

### Western blots

Proteins were resuspended in Laemli 1X, boiled for 5 minutes, separated by 10% SDS-PAGE and transferred to a nitrocellulose membrane (Protran). Membranes were blocked with 5% non-fat milk (weight/volume) in TBST (TBS with 0.5% Tween-20) and incubated in 1% non-fat milk in TBST with the primary antibodies overnight at 4°C followed by three times 5 min washes in TBS at RT, and incubation with secondary antibodies conjugated with horseradish peroxidase (HRP) for 1h at RT, and finally washed three times 10 min in TBST at RT. Enzymatic activity was detected using the SuperSignal West Pico/Femto Chemiluminescent Substrate (Genetex) or Immobilon (Roche). The primary antibodies used were: anti-RPAP3 (Sigma SAB1411438-100UG, 1/1000 dilution), anti-RUVBL1 (Proteintech 10210-2-AP, 1/500 dilution), anti-GAPDH (Abcam 8245, 1/20000 dilution) and anti-GFP (Roche 11814460001, 1/1000 dilution). Secondary antibodies were anti-rabbit (Abcam ab6721, dilution 1/10000) and anti-mouse (Abcam ab6728, dilution 1/5000) coupled to HRP.

### Immunofluorescence

Cells grown on coverslips were washed with PBS and fixed with 4% paraformaldehyde in PBS for 20 min at RT. Permeabilization was done in 0.1 % Triton X-100 in PBS for 10 min at RT. Coverslips were then incubated in PBS/BSA 1% with primary antibodies for 1h at RT, washed twice in PBS, incubated with secondary antibody for 1h at RT and washed again twice 10 min at RT. Primary antibodies used were anti-POLR2A (PB-7C2, Euromedex, 1/100 dilution) and anti-RPAP3 (Sigma SAB1411438-100UG, 1/100 dilution). Secondary antibodies were anti-mouse antibodies (Jackson 15-165-003, 1/1000 dilution) or anti-rabbit (Jackson 111-166-047, 1/200 dilution) coupled to Cyanine 3. Coverslips were mounted in Vectashield medium containing DAPI (Vector Laboratories, Inc.).

### SmiFISH with DNA probes

Cells were grown on glass coverslips or glass-bottom 96-well plates (Greiner 655892), and fixed with 4% paraformaldehyde in PBS for 20 min at RT and permeabilized with 70% ethanol overnight at 4 °C. To detect a specific mRNA, 24–48 unlabeled primary DNA probes complementary to the RNA of interest were used (Table S3). In addition to hybridizing to their targets, these probes contained a FLAP sequence that was pre-hybridized to a complementary secondary probe coupled to a fluorophore, as described previously ^71^. Pre-hybridization was done in NEBuffer 3 (NEB, 10 mM NaCl, 5 mM Tris-HCl, 1 mM MgCl_2_, pH 7.9) according to the ratios indicated in Table S3 and using a thermocycler with the following program: 85 °C for 3 min, 65 °C for 3 min, and 25 °C for 5 min. Cells on coverslips were first incubated in 1x SSC, 5M urea (Sigma-Aldrich) for 15 min at RT, and then incubated with the hybridization mixture containing the probe duplexes (2 µl of pre-hybridization mix per 100 µl of final volume), with 1X SSC, 0.34 mg/ml tRNA, 5M urea, 2 mM VRC, 0.2 mg/ml RNAse-free BSA, 10% Dextran sulfate, overnight in a wet chamber at 37°C. The day after, the samples were washed three times for 30 min in the wash buffer (1xSSC, 5M urea) at 37 °C and rinsed twice in PBS at RT. The 96-plates were incubated in the dark with 5µg/mL DAPI for 40 minutes at RT, then mounted in Vectashield medium without DAPI (Vector Laboratories, Inc.).

### SmFISH in 96-well plates with RNA probes

Cells were grown in a 96-well plate with glass bottom (Greiner 655892), fixed with 4% paraformaldehyde in PBS at RT for 20 minutes and permeabilized with 70% ethanol overnight at 4 °C. Probe sets of DNA oligonucleotides (GenScript) targeting the mRNAs of interest were generated based on the Oligostan script ^71^. Each oligo is composed of a specific hybridization sequence flanked by 2 readout sequences and a barcode shared among the gene probeset (Table S3). Gene specific probes were amplified by PCR with the gene-specific barcodes before being transcribed *in vitro* to obtain transcript-specific primary RNA probesets, as described previously ^72^. For each mRNA of interest, 50 ng of primary RNA probes and 25 ng of each fluorescent secondary DNA probes (TYE-650-labeled LNA/DNA oligonucleotides, Qiagen, ^72^ targeting the readout sequences were pre-hybridized in 100µl of a solution containing 7.5 M urea (Sigma-Aldrich), 0.34 mg/ml tRNA (Sigma-Aldrich) and 10% Dextran sulfate. Pre-hybridization was performed in a thermocycler, with the following program: 90 °C for 3 min, 53 °C for 15 min. Cells in 96-well plates were washed with PBS and incubated 15 min at RT in the hybridization buffer (1xSSC, 7.5 M urea) before adding 100µl of the hybridization mixture in each well. Hybridization was performed at 48°C overnight. The next day, the plate was washed 8 times for 20 min each in 1xSSC, and 7.5 M urea at 48°C. Finally, cells were washed two times with PBS at RT for 10 min each, stained with 5 µg/mL DAPI, and mounted in 90% glycerol (VWR), 1 mg/mL p-Phenylenediamine (Sigma-Aldrich), in PBS pH 8.

### Image acquisition

Coverslips images were acquired using a 63x oil immersion objective (1.4 NA, PL Apo, oil) on a Leica DM6B microscope. DAPI, GFP, CY3 and CY5 channels were imaged using the following filters combination : 405/60, 455, 470/ 40; 470/40, 495, 525/50; 545/30, 570, 610/75; 620/60, 660, 700/75. Alternatively, coverslips were imaged using a 63× objective (Plan Apochromat, 1.4 NA, oil) on a Zeiss Axioimager Z2 widefield microscope. DAPI, GFP, CY3 and CY5 channels were then imaged using the following filters combination: <365, FT 395, 445/50; 470/40, FT 495, 525/50; 545/25, 570, 605/70; 610/30, 660, 640/50. Images were taken as Z-stacks with one plane every 0.24 µm (Zeiss Z2) or every 0.376 µm (Leica Thunder). 96-well plates were imaged on an Opera Phenix High-Content Screening System (PerkinElmer), with a 63× water-immersion objective (NA 1.15). Three-dimensional images were acquired, consisting of 18 slices with a spacing of 0.6 µm. Figures were constructed using Omero, ImageJ and Adobe Illustrator.

### Image analysis

For Figure 5B, 6C and 8B, image quantifications were based on an image analysis pipeline that included cell segmentation, threshold-based spot detection, dense region deconvolution and a Density Based Spatial Clustering of Application with Noise (DBSCAN) algorithm. Firstly, nuclei and cytoplasm segmentation was inferred in 2D on image stack mean projection using Cellpose neural network ^73^. Secondly, single molecules were detected as 3D spots using the Big-FISH package ^74^ which relies on an intensity Laplacian of Gaussian (LoG) filter followed by a local maxima filter and a user-adjusted threshold. During analysis, we ensured that the same thresholds were applied on images made for the same RNA, for all cell treatments. As local maxima detection can lead to a misestimation of single molecule number in bright regions where individual molecules cannot be distinguished, a dense region deconvolution step was added. It consists of fitting the median detected spot with a 3D-gaussian to construct a reference spot and then reconstructing the bright region signal using multiple reference spots, thus providing the number of single RNAs in bright regions. Thirdly, a DBSCAN was used to assign spots to clusters (foci). At this point detected spots were divided in clustered spots (spots assigned to a cluster) or free spots. Finally, a centroid calculation was performed to assign coordinates to each cluster. This single molecule quantification process was repeated on all smFISH or SunTag channels, enabling quantification of RNA colocalization. During analysis, filtering was used to discard aberrations: (i) cells without nucleus or with more than one nuclei; (ii) any cell whose mask laid on the border of the field of view and that was thus considered incomplete; (iii) any spot detected outside of segmented cells; (iv) any regions containing an abnormally high average of 2 spots per pixel in a sphere of 1400nm. Distances between spots were computed using the exact Euclidian distance. Finally, spots were considered colocalizing when found closer together than a cutoff distance of 310 nm.

For the other Figures, a pipeline named SmallFish was developed to quantify single mRNAs and clusters, and evaluate the percentage of colocalization between clusters. It provides a ready to use graphical interface to combine the python packages used for cell analysis. In the pipeline, cell segmentation (2D) was performed using Cellpose (https://github.com/MouseLand/cellpose), while spot detection was carried out using BigFish (https://github.com/fish-quant/big-fish). Clusters were considered colocalizing when found closer together than a cutoff distance of 310 nm. Boxplots graphs were constructed on Pyhton using Seaborn package ^75^ and Statannotations package ^76^.

### RNA immunoprecipitation for RIP-Seq

HCT-116 cells with GFP-tagged RUVBL1 or RPAP3 alleles were seeded in 15 cm plates (1 plate/condition). At 70% confluency, cells were left untreated or treated as follow: with puromycin for 30 min at 100 μg/ml; with CB-6644 for 16h at 500 nM; with both CB-6644 for 16h and puromycin for 30 min. All treatments were done at 37°C. After treatments, cells were put on ice and washed twice with ice-cold PBS, and then lysed with 0.5 ml HNTG lysis buffer (20 mM Hepes pH 7.9, 150 mM NaCl, 1% triton X100, 10% glycerol, 1 mM MgCl_2_ and 1 mM EGTA) supplemented with a cocktail of protease inhibitors (Roche). Cells were incubated for 30 min on a tube roller at 4°C before being centrifuged for 10 min at 16000g at 4°C. The supernatant was subjected to immunoprecipitation for 1.5h at 4°C using GFP-Trap agarose beads (Chromotek-gta-20). Agarose control beads (Chromotek-bab-20) were used for non-specific control IP. All experiments were done in duplicates. Beads were washed 5 times with ice-cold HNTG. RNeasy mini kit (Qiagen-74106) was used to purify RNAs. DNA contamination was eliminated by a TURBO-DNA Free kit, as recommended by the manufacturer (Invitrogen-AM1907).

### RNA-sequencing and analysis

SMART-Seq Stranded Kit (Takara Bio) was used to generate cDNA libraries for high throughput sequencing using NovaSeq 6000 Illumina. RNA and DNA libraries quality were subjected to quality control checks periodically (or after each step) using a fragment analyzer (Kit high sensivity NGS) and qPCR (Roche Light Cycler 480). Ribosomal cDNAs were cleaved with scZapR enzyme in the presence of scR-Probes designed against mammalian ribosomal RNAs. Libraries were then sequenced with NovaSeq Reagent Kits on a flow cell SP (single read −100 nucleotides) using the 2-channel sequence by synthesis method. Before demultiplexing, sequences were mixed with PhiX sequences as an Illumina control (not indexed). Demultiplexing was realized using Illumina bcl2fastq software (v2.20.0.422) and all PhiX sequences were removed.

FastaQC software (v0.11.9) provided a tool to check read quality after high-throughput sequencing and MultiQC program (V1.12) served to group data that emerges from all samples. FastQ screen software served to detect contaminating genomes. Sequences were aligned on 15 species genomes, mycoplasma, and ribosomal RNAs using Bowtie2 software. No contamination was detected.

TopHat2 software (v2.1.1) was used to align the obtained reads on the human genome (hg38 version) and to annotate them according to NBCI database. FeatureCounts software was utilized to count reads per gene. All reads that overlap on two genes or show alignment at different positions on the genome were discarded. Counts were normalized in each sample to eliminate differences due to manipulation. This was done by a normalization method known as relative log expression (RLE). Differential expression genes were identified using EdgeR (v3.34.1). Tests are based on a generalized linear model and allow counts modeling using a negative binomial distribution. To calculate the reads per kilobase per million mapped reads (RPKM), the formula below was used: (read counts aligned on a specific gene/ (total aligned counts x gene length) x109.

### Label-free quantitative proteomics

HCT-116 cells with RUVBL1 or RPAP3 endogenously tagged with GFP were seeded in 15 cm plates (1 plate/condition). At 70% confluency, cells were treated or not with CB-6644 for 16h at 500 nM. Cells were washed with ice-cold PBS, then lysed with 0.5 ml HNTG lysis buffer supplemented with proteinase inhibitor cocktail (Roche; see above for the composition of HNTG). Cells were scraped and incubated for 30 min at 4°C before being centrifuged for 10 min at 16000g at 4°C. The supernatant was subjected to immunoprecipitation using GFP-Trap agarose beads (Chromotek-gta-20). Agarose control beads (Chromotek-bab-20) were used for non-specific control IP. All experiments were done in triplicates. Extracts were incubated with beads for 1.5h at 4°C and washed 5 times with ice-cold HNTG. Proteins were then eluted in Laemmli buffer and heated 10 min at 95°C.

For sample digestion, proteins were loaded onto SDS-PAGE (10 % polyacrylamide, Mini-PROTEAN® TGX™ Precast Gels, Bio-Rad, Hercules USA) and stained with Protein Staining Solution (Euromedex, Ref 10-0911). Gel lanes were cut into one single band and destained with 50 mM TriEthylAmmonium BiCarbonate (TEABC, Merck, Ref T7408) followed by three washes in 100% acetonitrile. After reduction (with 10 mM dithiothreitol in 50 mM TEABC at 60 °C for 30 min) and alkylation (with 55 mM iodoacetamide in 50 mM TEABC at room temperature for 30 min), proteins were digested in-gel using trypsin (1 µg/band, Gold, Promega, Ref V5280). Digest products were vacuum-dried before LC MS/MS analysis. For mass spectrometry, a Q Exactive HF mass spectrometer (Thermo Fisher Scientific) coupled with an Ultimate 3000 nano-HPLC (Thermo Fisher Scientific) was used. Peptide samples were solubilized in 0.05% trifluoroacetic acid (TFA) / 2% ACN, and were injected for desalting and pre-concentration on a PepMap®100 C18 precolumn (0.3 mm x 5 mm; ThermoFisher Scientific). Peptides separation were done on a 50 cm analytical reversed-phase column (75 mm inner diameter, Acclaim Pepmap 100® C18, Thermo Fisher Scientific) using a 120 min gradient of 2 to 40% of buffer B (80% ACN, 0.1% formic acid) and a flow rate of 300 nL/min. MS/MS analyses were performed in a data-dependent mode. Full scans (375-1500 m/z) were acquired on the Orbitrap mass analyzer with a 60 000 resolution at 200 m/z. For the full scans, 3e^6^ ions were accumulated within a maximum injection time of 60 ms and detected in the Orbitrap analyzer. The twelve most intense ions with charge states ≥ 2 were sequentially isolated to a target value of 1e^5^ with a maximum injection time of 100 ms and fragmented by HCD (Higher-energy collisional dissociation) in the collision cell (normalized collision energy of 28%) and detected in the Orbitrap analyzer at 30 000 resolution. First mass was set to 100 m/z.

Raw spectra were processed using the MaxQuant environment (v.2.0.3.0), and Andromeda for database search with label-free quantification (LFQ), match between runs and the iBAQ algorithm (CIT). The MS/MS spectra were matched against the UniProt Reference proteome of Homo sapiens (Proteome ID UP000005640; release 2022_03; https://www.uniprot.org/) and a homemade contaminant database. Different release versions were used, depending on the date of analysis. Enzyme specificity was set to trypsin/P, and the search included cysteine carbamidomethylation as a fixed modification and oxidation of methionine, and acetylation (protein N-term) as variable modifications. Up to two missed cleavages were allowed for protease digestion. FDR was set at 0.01 for peptides and proteins and the minimal peptide length at 7. A representative protein ID in each protein group was automatically selected using an in-house developed bioinformatics tool (Leading tool 3.4) as the best annotated protein in UniProtKB (reviewed entries rather than automatic ones ^77^. Analysis of quantification data, statistical analyses and graphical representations were performed using Perseus (v1.6.15).

## Acknowledgements

We thank Drs. D. Helmlinger and B. Pradet-Balade for stimulating discussions and advices, and Dr. B. Delaval for the gift of the Hela centrin-GFP cell line. We acknowledge (i) the sequencing facility MGX, which obtained financial support from France Génomique National infrastructure, funded as part of the “Investissement d’Avenir” program (contract ANR-10-INBS-09); (ii) the microscopy facility MRI, member of the national infrastructure France-BioImaging supported by the French National Research Agency (ANR-10-INBS-04); and (iii) Montpellier Proteomics Platform (PPM, BioCampus Montpellier) supported by the French National Research Agency via ProFI fundings (Proteomics French Infrastructure, ANR-24-INBS-0015). The work was supported by grants from the ANR (ANR-23-CE12-0022), the INCa (PLBIO23) and the ‘Ligue Nationale contre le Cancer’ (équipe labélisée).

## Author contributions

EB conceived the study and EB, CV and SB supervised the work. SS optimized the RIP-seq and performed the initial experiment of Figure 1. MP, with help of HC, SB and CV, performed the other experiments and interpreted the data. FS developed small-FISH and quantified the microscopy images. JI and SG performed the RNA sequencing and analyzed the data. SU and MS did the mass spectrometer analysis and processed the data. EB, CV and MP wrote the manuscript, which was corrected by all the authors.

## Supplementary Figures

**Figure S1. Characterization of HCT116 RUVBL1-GFP cells RIP-seq analyses**

**A-** Schematic of the PCR genotyping of the cassette integrated at the RUVBL1 genomic locus. PCR products are indicated in red below the cassette (1 and 2). The gel image shows the results of PCR genotyping. The size of the DNA ladder is indicated on the left.

**B-** Western blot with anti-RUVBL1 and anti-GAPDH antibodies on extracts from parental and RUVBL1-GFP HCT116 cells. The size of the protein ladder is indicated on the left. The names of the detected proteins are on the right.

**C-** The graph shows the enrichment of the mRNAs found in the GFP IP over the control IP, in puromycin treated cells (y axis) versus untreated cells (x axis). Each dot represents a mRNA and is colored according to the code of Figure 1. Only the top 200 mRNAs with the best p-values in each IP are shown. **D-** Same as in C but with cells treated with CB-6644 instead of puromycin.

**Figure S2. Characterization of HCT116 RPAP3-GFP cells and RIP-Seq analyses**

**A-** Scheme of the PCR genotyping for the cassette integrated at the RPAP3 genomic locus. PCR products are indicated in red below the cassette (1 and 2). The gel image shows the genotyping results. The size of the DNA ladder is indicated on the left.

**B-** Western blot with anti-RPAP3 and anti-GAPDH antibodies on extracts from parental and RPAP3-GFP HCT116 cells. The size of the protein ladder is indicated on the left. The name of the detected proteins are on the right.

**C-** The graph shows the enrichment of the mRNAs found in the GFP IP over the control IP, for puromycin treated cells (y axis) and untreated cells (x axis). Each dot represents a mRNA and is color coded according as in Figure 3. Only the top 200 mRNAs with the best p-value in each IP are shown.

**Figure S3. mRNAs enriched in the RIP-Seq experiments are recruited to DsRed2-DDX6fl-RUVBL1 artificial condensates**

Micrographs of HeLa cells stably expressing DsRed2-DDX6fm-RUVBL1 (middle) and labelled for the indicated mRNAs by smFISH (left). Merge images (right) show mRNAs in red, DsRed2-DDX6fm-RUVBL1 in green and DAPI staining in blue. Scale bar is 10 µm. Arrowheads point to the artificial RUVBL1 condensates. Inset shows a higher magnification of the bowed area.

**Figure S4. Characterization of HeLa RPAP3-SunTag*32 cells and image quantifications**

**A-** Scheme for the PCR genotyping of the cassette integrated at the RPAP3 genomic locus. PCR products are indicated in red below the cassette (1 and 2). The gel image shows the genotyping results. The size of the DNA ladder is indicated on the left.

**B-** Western blot with anti-RPAP3 and anti-GAPDH antibodies on extract from parental and RPAP3-SunTag*32 cells. The size of the protein ladders is indicated on the left. The name of the detected proteins are on the right. The panel on the left shows a high exposure of the signal for the RPAP3-SunTag*32 cells.

**C-** Quantification of the colocalization between single molecules of mRNAs and RPAP3-SunTag. Boxplots represent the colocalization between single POLR2A (left) or PRPF8 (right) mRNAs, and single molecules of RPAP3-SunTag in normal (unt, blue) or treated (puro, light blue) conditions. Each observation is a cell, at least 200 cells were analyzed. P-values were determined using Welch’s t-test. * P<0.05, ** P<0.01, *** P<0.001, *** P<0.0001.

**D-** Quantification of the number of endogenous RPAP3 foci in HeLa Kyoto cells, in untreated (unt, blue) or puromycin (puro, light blue) treated cells. Boxplots represent the number of endogenous RPAP3 foci per cell in both conditions. Each observation is a cell, at least 200 cells were analyzed. P-values were determined using Welch’s t-test. * P<0.05, ** P<0.01, *** P<0.001, *** P<0.0001.

**E-** Quantification of the number of mRNA foci in HeLa Kyoto cells treated or not with puromycin. Boxplots represent the number of POLR2A (left), PRPF8 (middle) and DYNC1H1 (right) mRNA foci in cells untreated (unt, blue) or treated with puromycin (puro, light blue) cells. Each observation is a cell, at least 125 cells were analyzed. P-values were determined using Welch’s t-test. * P<0.05, ** P<0.01, *** P<0.001, *** P<0.0001.

**F-** Micrographs of HeLa Kyoto cells labelled for POLR2A (left) and DYNC1H1 (middle) mRNAs by dual color smFISH. Merge images (right) show POLR2A mRNA in red, DYNC1H1 in green and DAPI staining in blue. Scale bar is 10 µm. Red arrowheads point to POLR2A mRNA foci and green arrowheads point to DYNC1H1 mRNA foci.

**G-** Micrographs of HeLa Kyoto cells in normal or puromycin treated conditions and labelled for POLR2A mRNA (left) by smFISH and for POLR2A protein (middle) by immunofluorescence. Merge images (right) show POLR2A mRNA in red, POLR2A protein in green and DAPI staining in blue. Scale bar is 10µm. Arrowheads point to foci or mRNA and protein. Inset shows a higher magnification of the boxed area.

**Figure S5. Dual color smFISH of POLR1A and POLR2A mRNAs in HeLa cells**

**A-** Micrographs of HeLa cells labelled by dual color smFISH for POLR2A (left) and POLR1A (middle) mRNAs. Merge images (right) show POLR2A mRNA in red, POLR1A mRNA in green and DAPI staining in blue. Scale bars are 20µm.

**B-** Quantification of the colocalization of POLR2A with POLR1A mRNAs, and with the negative control JUP mRNA. Boxplots represent the proportion of colocalization (y axis, %) between pairs of mRNAs under normal (unt, blue) and puromycin conditions (puro, light blue). Each observation is a cell, at least 200. P-values were determined using Welch’s t-test. * P<0.05, ** P<0.01, *** P<0.001, *** P<0.0001.

## References

1. Broeck, A. V. & Klinge, S. Eukaryotic Ribosome Assembly.

2. Massenet, S., Bertrand, E. & Verheggen, C. Assembly and trafficking of box C/D and H/ACA snoRNPs. RNA Biology 14, 680–692 (2017).

3. Lanfranco, M., Vassallo, N. & Cauchi, R. J. Spinal Muscular Atrophy: From Defective Chaperoning of snRNP Assembly to Neuromuscular Dysfunction. Front Mol Biosci 4, 41 (2017).

4. Garrido-Godino, A. I., Gutiérrez-Santiago, F. & Navarro, F. Biogenesis of RNA Polymerases in Yeast. Frontiers in Molecular Biosciences 8, (2021).

5. von Morgen, P., Hořejší, Z. & Macurek, L. Substrate recognition and function of the R2TP complex in response to cellular stress. Front. Genet. 6, (2015).

6. Pal, M., et al. Structure of the TELO2-TTI1-TTI2 complex and its function in TOR recruitment to the R2TP chaperone. Cell Reports 36, (2021).

7. Serna, M. et al. CryoEM of RUVBL1-RUVBL2-ZNHIT2, a complex that interacts with pre-mRNA-processing-splicing factor 8. Nucleic Acids Res 50, 1128–1146 (2022).

8. Lynham, J. & Houry, W. A. The Role of Hsp90-R2TP in Macromolecular Complex Assembly and Stabilization. Biomolecules 12, 1045 (2022).

9. Boulon, S. et al. The Hsp90 chaperone controls the biogenesis of L7Ae RNPs through conserved machinery. The Journal of Cell Biology 180, 579–595 (2008).

10. Zhao, R. et al. Molecular chaperone Hsp90 stabilizes Pih1/Nop17 to maintain R2TP complex activity that regulates snoRNA accumulation. J Cell Biol 180, 563–578 (2008).

11. Machado-Pinilla, R., Liger, D., Leulliot, N. & Meier, U. T. Mechanism of the AAA+ ATPases pontin and reptin in the biogenesis of H/ACA RNPs. RNA 18, 1833–1845 (2012).

12. Venteicher, A. S., Meng, Z., Mason, P. J., Veenstra, T. D. & Artandi, S. E. Identification of ATPases pontin and reptin as telomerase components essential for holoenzyme assembly. Cell 132, 945–957 (2008).

13. Malinová, A. et al. Assembly of the U5 snRNP component PRPF8 is controlled by the HSP90/R2TP chaperones. Journal of Cell Biology 216, 1579 (2017).

14. Cloutier, P. et al. R2TP/Prefoldin-like component RUVBL1/RUVBL2 directly interacts with ZNHIT2 to regulate assembly of U5 small nuclear ribonucleoprotein. Nature Communications 8, 1–14 (2017).

15. Abel, Y. et al. The interaction between RPAP3 and TRBP reveals a possible involvement of the HSP90/R2TP chaperone complex in the regulation of miRNA activity. Nucleic Acids Research 50, 2172–2189 (2022).

16. Boulon, S. et al. HSP90 and Its R2TP/Prefoldin-like Cochaperone Are Involved in the Cytoplasmic Assembly of RNA Polymerase II. Molecular Cell 39, 912–924 (2010).

17. Zur Lage, P., et al. Ciliary dynein motor preassembly is regulated by Wdr92 in association with HSP90 co-chaperone, R2TP. J Cell Biol 217, 2583–2598 (2018).

18. Horejsí, Z. et al. CK2 phospho-dependent binding of R2TP complex to TEL2 is essential for mTOR and SMG1 stability. Mol Cell 39, 839–850 (2010).

19. Takai, H., Xie, Y., de Lange, T. & Pavletich, N. P. Tel2 structure and function in the Hsp90-dependent maturation of mTOR and ATR complexes. Genes Dev 24, 2019–2030 (2010).

20. Abéza, C. et al. The HSP90/R2TP Quaternary Chaperone Scaffolds Assembly of the TSC Complex. Journal of Molecular Biology 436, 168840 (2024).

21. Sardiu, M. E. et al. Probabilistic assembly of human protein interaction networks from label-free quantitative proteomics. Proc Natl Acad Sci U S A 105, 1454–1459 (2008).

22. Cloutier, P. et al. High-resolution mapping of the protein interaction network for the human transcription machinery and affinity purification of RNA polymerase II-associated complexes. Methods 48, 381–386 (2009).

23. Seraphim, T. V. et al. Assembly principles of the human R2TP chaperone complex reveal the presence of R2T and R2P complexes. Structure 30, 156–171.e12 (2022).

24. Bizarro, J. et al. Proteomic and 3D structure analyses highlight the C/D box snoRNP assembly mechanism and its control. Journal of Cell Biology 207, 463–480 (2014).

25. Maurizy, C. et al. The HSP90/R2TP assembly chaperone promotes cell proliferation in the intestinal epithelium. Nat Commun 12, 4810 (2021).

26. Maurizy, C. et al. The RPAP3-Cterminal domain identifies R2TP-like quaternary chaperones. Nature Communications 9, 1–16 (2018).

27. Zhao, R. et al. Navigating the Chaperone Network: An Integrative Map of Physical and Genetic Interactions Mediated by the Hsp90 Chaperone. Cell 120, 715–727 (2005).

28. Houry, W. A., Bertrand, E. & Coulombe, B. The PAQosome, an R2TP-Based Chaperone for Quaternary Structure Formation. Trends Biochem Sci 43, 4–9 (2018).

29. Muñoz-Hernández, H. et al. Structural mechanism for regulation of the AAA-ATPases RUVBL1-RUVBL2 in the R2TP co-chaperone revealed by cryo-EM. Science Advances 5, eaaw1616 (2019).

30. Gorynia, S. et al. Structural and functional insights into a dodecameric molecular machine - the RuvBL1/RuvBL2 complex. J Struct Biol 176, 279–291 (2011).

31. Zhou, C. Y. et al. Regulation of Rvb1/Rvb2 by a Domain within the INO80 Chromatin Remodeling Complex Implicates the Yeast Rvbs as Protein Assembly Chaperones. Cell Rep 19, 2033–2044 (2017).

32. Martino, F. et al. RPAP3 provides a flexible scaffold for coupling HSP90 to the human R2TP co-chaperone complex. Nature Communications 9, 1–13 (2018).

33. Silva, S. T. N. et al. X-ray structure of full-length human RuvB-Like 2 - mechanistic insights into coupling between ATP binding and mechanical action. Sci Rep 8, 13726 (2018).

34. McKeegan, K. S., Debieux, C. M., Boulon, S., Bertrand, E. & Watkins, N. J. A dynamic scaffold of pre-snoRNP factors facilitates human box C/D snoRNP assembly. Mol Cell Biol 27, 6782–6793 (2007).

35. Dauden, M. I., López-Perrote, A. & Llorca, O. RUVBL1–RUVBL2 AAA-ATPase: a versatile scaffold for multiple complexes and functions. Current Opinion in Structural Biology 67, 78–85 (2021).

36. Henri, J. et al. Deep Structural Analysis of RPAP3 and PIH1D1, Two Components of the HSP90 Co-chaperone R2TP Complex. Structure 26, 1196–1209.e8 (2018).

37. Benbahouche, N. E. H. et al. Drosophila Spag is the homolog of RNA polymerase II-associated protein 3 (RPAP3) and recruits the heat shock proteins 70 and 90 (Hsp70 and Hsp90) during the assembly of cellular machineries. J Biol Chem 289, 6236–6247 (2014).

38. Kakihara, Y., Makhnevych, T., Zhao, L., Tang, W. & Houry, W. A. Nutritional status modulates box C/D snoRNP biogenesis by regulated subcellular relocalization of the R2TP complex. Genome Biol 15, 404 (2014).

39. Yu, G. et al. Yeast R2TP Interacts with Extended Termini of Client Protein Nop58p. Sci Rep 9, 20228 (2019).

40. Takai, H., Wang, R. C., Takai, K. K., Yang, H. & de Lange, T. Tel2 regulates the stability of PI3K-related protein kinases. Cell 131, 1248–1259 (2007).

41. Goto, G. H. et al. Two separate pathways regulate protein stability of ATM/ATR-related protein kinases Mec1 and Tel1 in budding yeast. PLOS Genetics 13, e1006873 (2017).

42. Hořejší, Z. et al. Phosphorylation-dependent PIH1D1 interactions define substrate specificity of the R2TP cochaperone complex. Cell Rep 7, 19–26 (2014).

43. Abel, Y. et al. NOPCHAP1 is a PAQosome cofactor that helps loading NOP58 on RUVBL1/2 during box C/D snoRNP biogenesis. Nucleic Acids Research 49, 1094–1113 (2021).

44. Peng, W. T. et al. A panoramic view of yeast noncoding RNA processing. Cell 113, 919–933 (2003).

45. Pal, M. et al. Structural Basis for Phosphorylation-Dependent Recruitment of Tel2 to Hsp90 by Pih1. Structure 22, 805–818 (2014).

46. Shiber, A. et al. Cotranslational assembly of protein complexes in eukaryotes revealed by ribosome profiling. Nature 561, 268–272 (2018).

47. Halbach, A. et al. Cotranslational assembly of the yeast SET1C histone methyltransferase complex. EMBO J 28, 2959–2970 (2009).

48. Bernardini, A. & Tora, L. Co-translational Assembly Pathways of Nuclear Multiprotein Complexes Involved in the Regulation of Gene Transcription. J Mol Biol 436, 168382 (2024).

49. Kamenova, I. et al. Co-translational assembly of mammalian nuclear multisubunit complexes. Nat Commun 10, 1740 (2019).

50. Bernardini, A. et al. Hierarchical TAF1-dependent co-translational assembly of the basal transcription factor TFIID. Nat Struct Mol Biol 30, 1141–1152 (2023).

51. Yayli, G. et al. ATAC and SAGA co-activator complexes utilize co-translational assembly, but their cellular localization properties and functions are distinct. Cell Rep 42, 113099 (2023).

52. Bertolini, M. et al. Interactions between nascent proteins translated by adjacent ribosomes drive homomer assembly. Science 371, 57–64 (2021).

53. Mallik, S. et al. Structural determinants of co-translational protein complex assembly. Cell 188, 764–777.e22 (2025).

54. Paknia, E., Chari, A., Stark, H. & Fischer, U. The Ribosome Cooperates with the Assembly Chaperone pICln to Initiate Formation of snRNPs. Cell Reports 16, 3103–3112 (2016).

55. King, T. H., Decatur, W. A., Bertrand, E., Maxwell, E. S. & Fournier, M. J. A Well-Connected and Conserved Nucleoplasmic Helicase Is Required for Production of Box C/D and H/ACA snoRNAs and Localization of snoRNP Proteins. Mol Cell Biol 21, 7731–7746 (2001).

56. Watkins, N. J. et al. Assembly and maturation of the U3 snoRNP in the nucleoplasm in a large dynamic multiprotein complex. Mol Cell 16, 789–798 (2004).

57. Assimon, V. A. et al. CB-6644 Is a Selective Inhibitor of the RUVBL1/2 Complex with Anticancer Activity. ACS Chem Biol 14, 236–244 (2019).

58. Yenerall, P. et al. RUVBL1/RUVBL2 ATPase Activity Drives PAQosome Maturation, DNA Replication and Radioresistance in Lung Cancer. Cell Chem Biol 27, 105–121.e14 (2020).

59. Dos Santos Morais, R., et al. Deciphering cellular and molecular determinants of human DPCD protein in complex with RUVBL1/RUVBL2 AAA-ATPases. Journal of Molecular Biology 434, 167760 (2022).

60. Safieddine, A. et al. A choreography of centrosomal mRNAs reveals a conserved localization mechanism involving active polysome transport. Nat Commun 12, 1352 (2021).

61. Jády, B. E., Bertrand, E. & Kiss, T. Human telomerase RNA and box H/ACA scaRNAs share a common Cajal body–specific localization signal. Journal of Cell Biology 164, 647–652 (2004).

62. Kittur, N., Darzacq, X., Roy, S., Singer, R. H. & Meier, U. T. Dynamic association and localization of human H/ACA RNP proteins. RNA 12, 2057–2062 (2006).

63. Tanenbaum, M. E., Gilbert, L. A., Qi, L. S., Weissman, J. S. & Vale, R. D. A protein-tagging system for signal amplification in gene expression and fluorescence imaging. Cell 159, 635–646 (2014).

64. Pichon, X. et al. Visualization of single endogenous polysomes reveals the dynamics of translation in live human cells. Journal of Cell Biology 214, 769–781 (2016).

65. Caudron-Herger, M., Jansen, R. E., Wassmer, E. & Diederichs, S. RBP2GO: a comprehensive pan-species database on RNA-binding proteins, their interactions and functions. Nucleic Acids Res 49, D425–D436 (2021).

66. Toullec, D. et al. The Hsp90 cochaperone TTT promotes cotranslational maturation of PIKKs prior to complex assembly. Cell Rep 37, 109867 (2021).

67. He, X. D. et al. Overlapping peri-implantation phenotypes of ZNHIT1 and ZNHIT2 despite distinct functions during early mouse development†. Biol Reprod 111, 1017–1029 (2024).

68. Schirle, N. T. & MacRae, I. J. The crystal structure of human Argonaute2. Science 336, 1037–1040 (2012).

69. Panasenko, O. O. et al. Co-translational assembly of proteasome subunits in NOT1-containing assemblysomes. Nat Struct Mol Biol 26, 110–120 (2019).

70. Rual, J.-F. et al. Human ORFeome version 1.1: a platform for reverse proteomics. Genome Res 14, 2128–2135 (2004).

71. Tsanov, N. et al. smiFISH and FISH-quant – a flexible single RNA detection approach with super-resolution capability. Nucleic Acids Res 44, e165–e165 (2016).

72. Safieddine, A. et al. HT-smFISH: a cost effective and flexible workflow for high-throughput single molecule RNA imaging. Nature Protocols 18, 157 (2023).

73. Pachitariu, M. & Stringer, C. Cellpose 2.0: how to train your own model. Nat Methods 19, 1634–1641 (2022).

74. Imbert, A. et al. FISH-quant v2: a scalable and modular tool for smFISH image analysis. RNA 28, 786–795 (2022).

75. Waskom, M. L. seaborn: statistical data visualization. Journal of Open Source Software 6, 3021 (2021).

76. Charlier, F., et al. trevismd/statannotations: v0.6. Zenodo 10.5281/zenodo.8396665 (2023).

77. Raynaud, F. et al. SNAP23-Kif5 complex controls mGlu1 receptor trafficking. J Mol Cell Biol 10, 423–436 (2018).

