## Supplementary figures and images for "Co-translational determination of quaternary structures in chaperone factories"

### Figures S1-S5

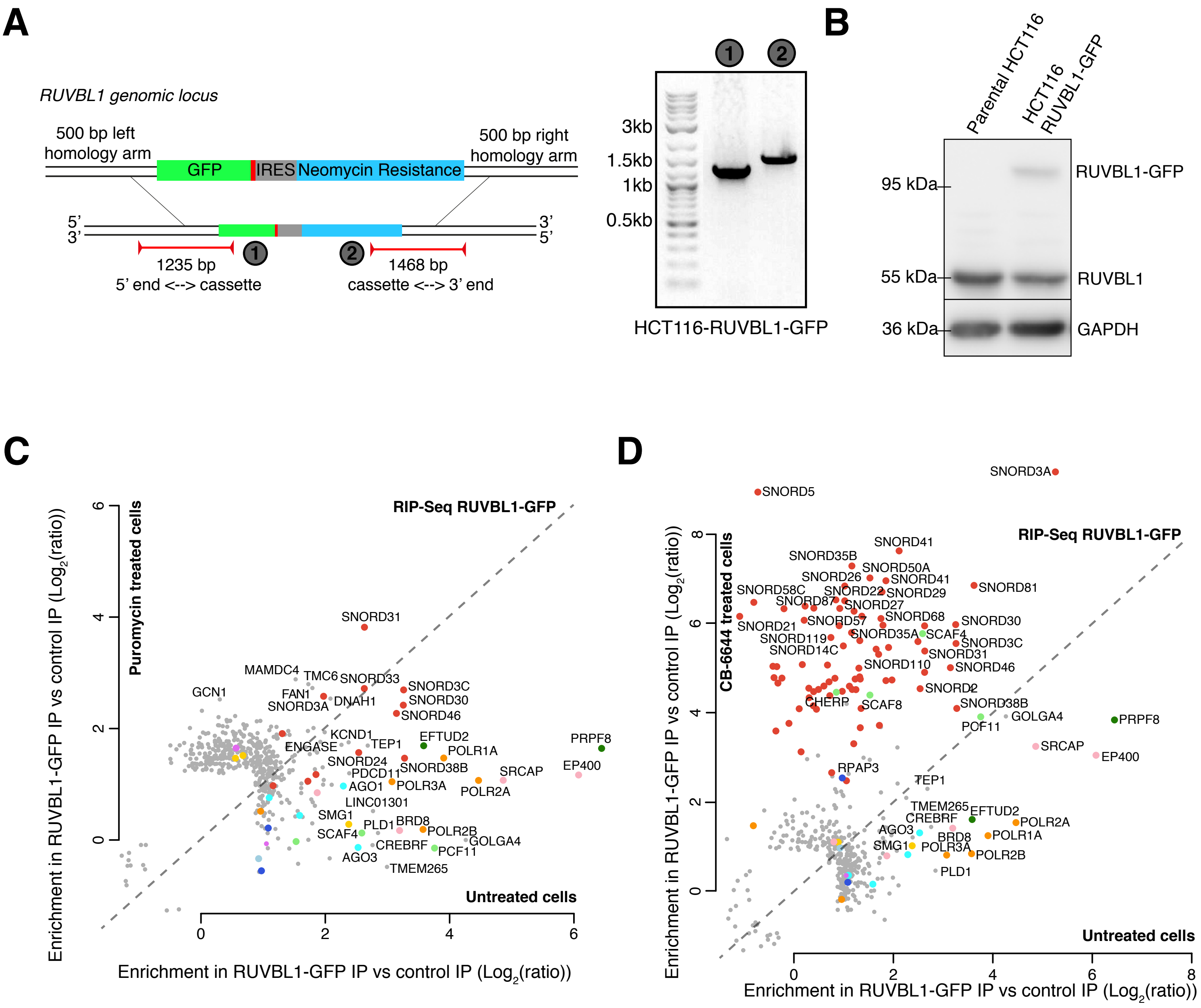

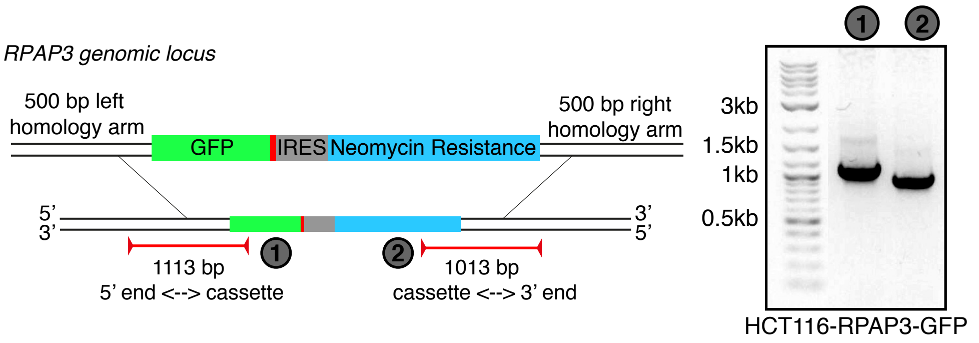

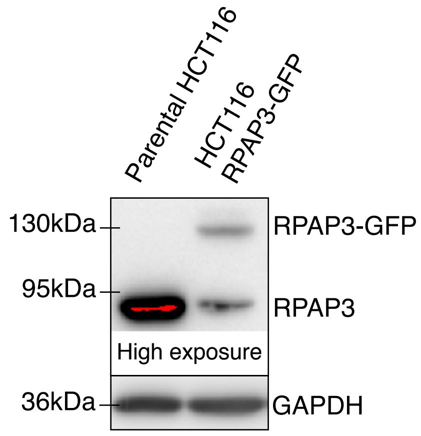

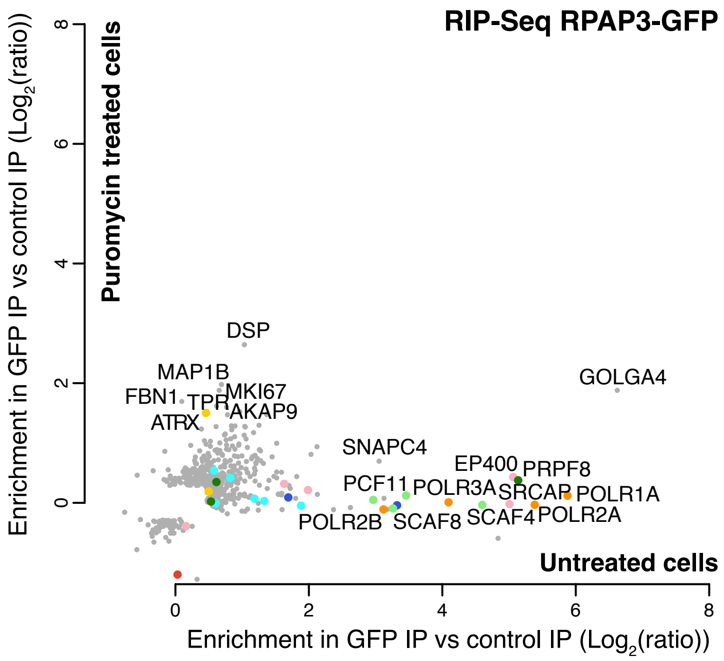

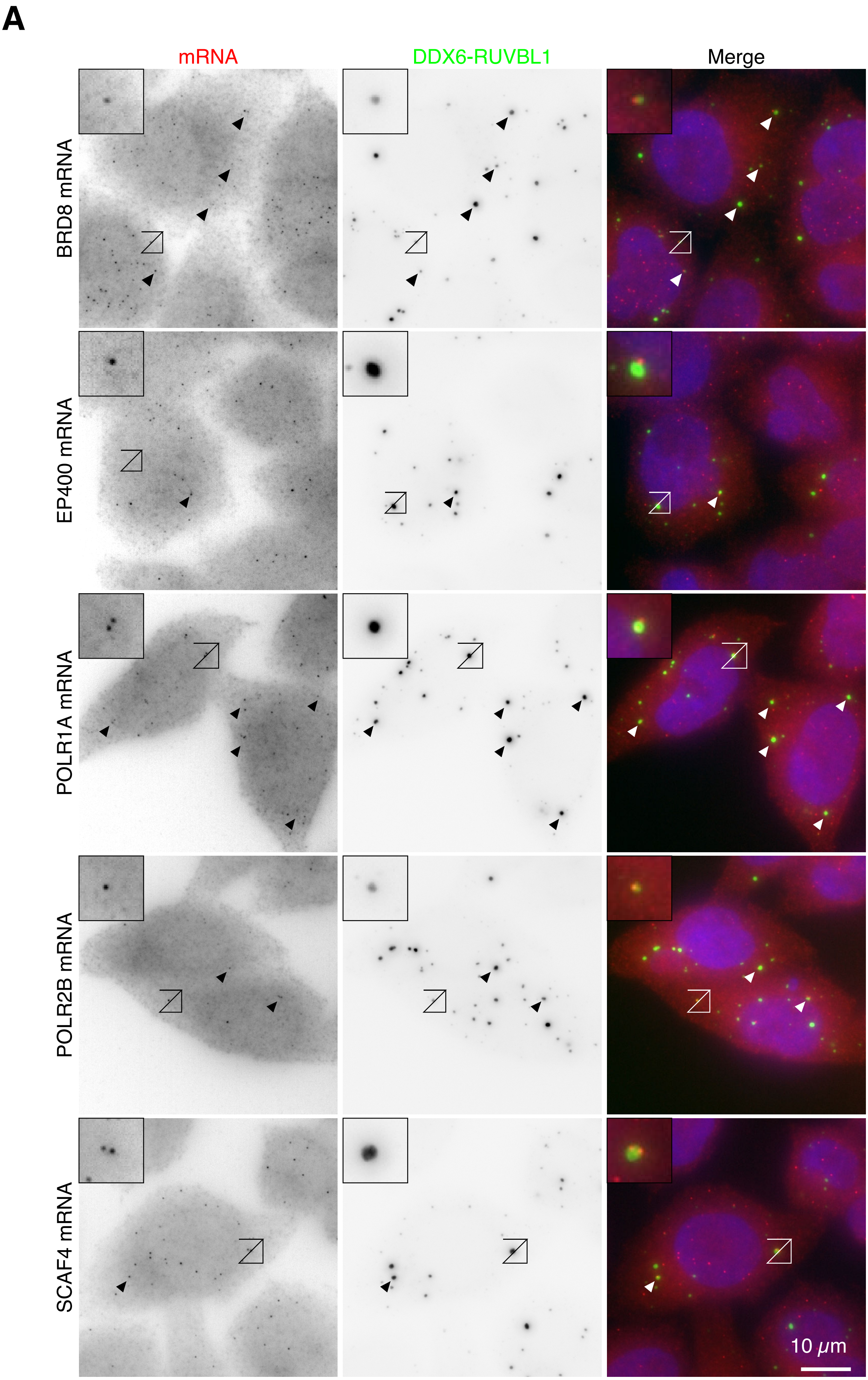

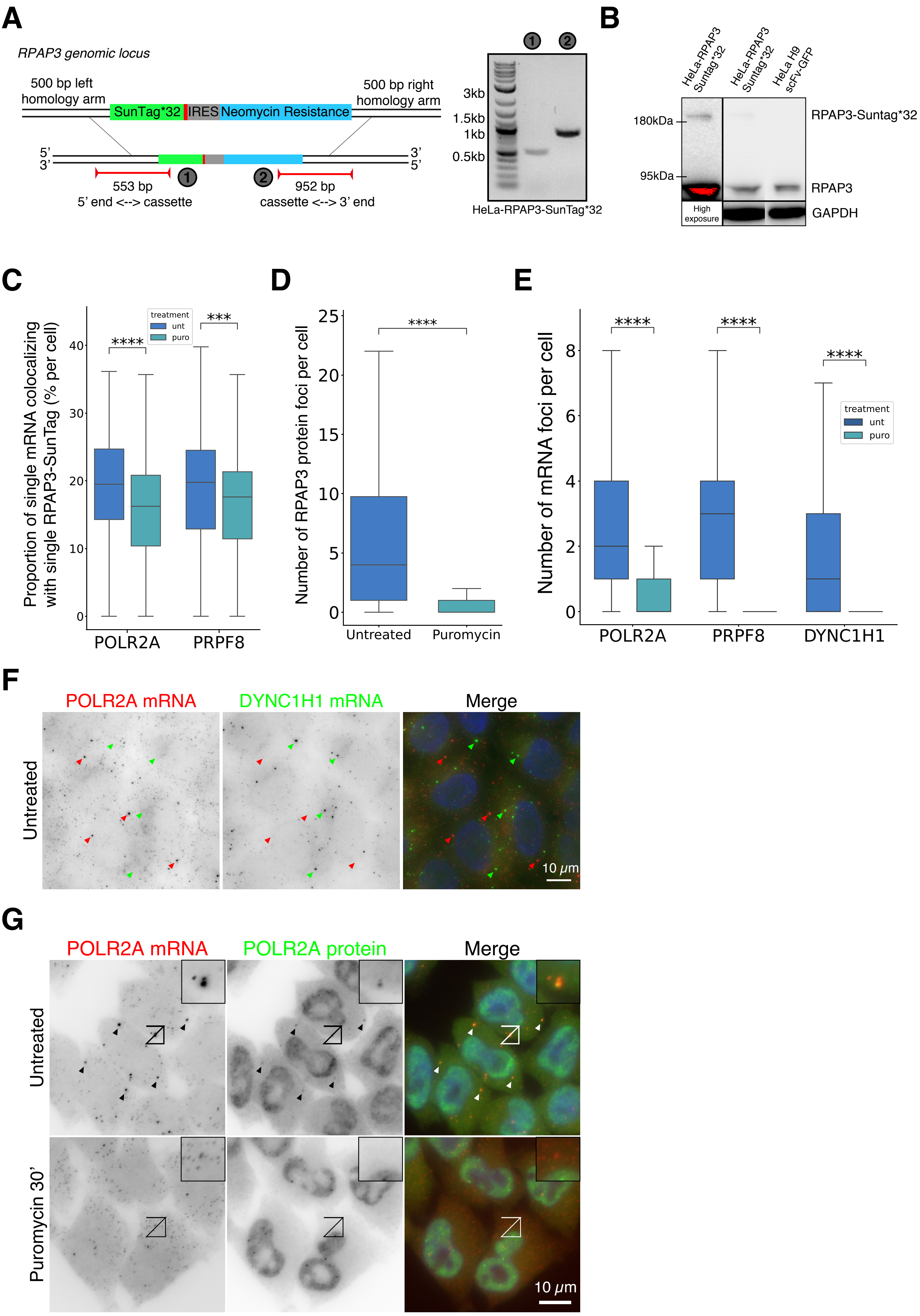

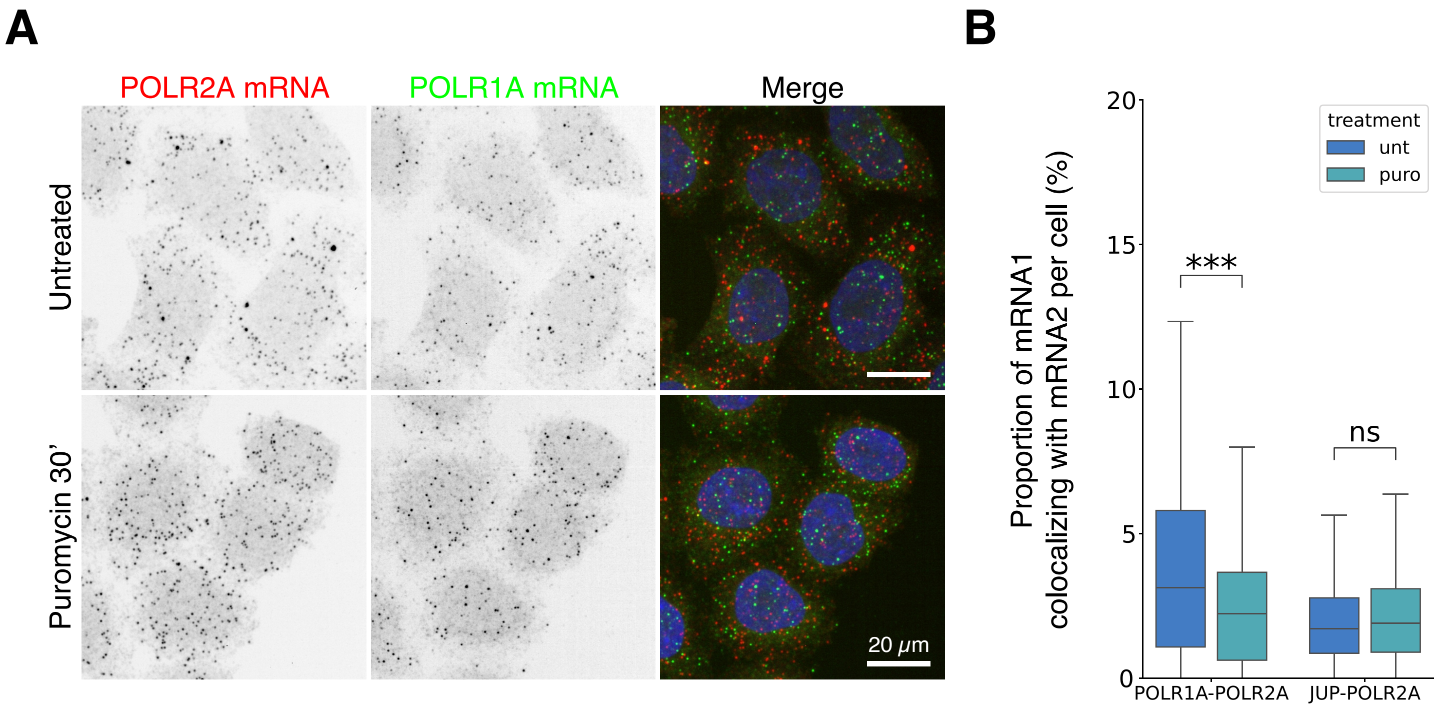
